# Mitochondrial dysfunction as a hallmark of brain senescence in telomerase-deficient mice

**DOI:** 10.64898/2026.08.25.746691

**Authors:** Debora Palomares, Joana Jorgji, Shirine Saleki, Tasha Ibrahim, Esther Paître, Axelle Loriot, Marc Dieu, Sophie Burteau, Patricia Renard, Manuel Johanns, Cyril Corbet, Laurent Gatto, Pascal Kienlen-Campard, Nuria Suelves

## Abstract

Neurodegenerative diseases, including Alzheimer’s disease (AD), are strongly associated with aging. However, the molecular mechanisms underlying pathological brain aging remain incompletely understood. In this study, we used a mouse model of telomere attrition, a major driver of cellular senescence, to perform an unbiased analysis of how telomere-driven senescence affects cellular physiology and contributes to processes relevant to neurodegenerative conditions.

After validating the presence of senescence hallmarks in telomerase-deficient brains, we characterized their transcriptomic and proteomic profiles. Mitochondrial function and associated energy metabolism emerged as the major dysregulated pathways, driven predominantly by proteomic rather than transcriptomic changes. Functional biochemical analyses on isolated brain mitochondria demonstrated impaired electron transport chain (ETC) complex activity and reduced energetic status, despite preserved ETC complex integrity and mitochondrial content. Further analyses in senescent primary neurons indicated an accumulation of dysfunctional mitochondria, characterized by increased reactive oxygen species (ROS) production and reduced ATP levels, although basal cellular respiration was maintained. At the tissue level, these alterations were associated with moderate reductions in neuronal density in the subiculum and cortical layer V, indicating region-specific vulnerability rather than widespread neurodegeneration.

We propose that a major consequence of telomere dysfunction associated with pathological brain aging is the downregulation of mitochondrial activity, which contributes to the selective vulnerability of specific brain regions. These findings highlight mitochondrial pathways as attractive targets for interventions aimed at preserving brain health during aging.

## 1 Introduction

Advances in medicine and public health over recent decades have markedly increased global life expectancy, yet they have also contributed to a rapidly aging population and an unprecedented rise in the prevalence of age-related diseases. Indeed, aging is recognized as the primary risk factor for many chronic conditions, including neurodegenerative diseases (J. Guo et al., 2022; Hou et al., 2019; S. Wang, Jiang, Yang, Meng, & Zhang, 2024). Millions of people suffer from neurodegenerative diseases, and despite their growing socioeconomic burden, effective and affordable treatments remain unavailable. Consequently, considerable research efforts have been directed towards understanding the biological mechanisms that drive aging and age-related neurodegeneration.

Cellular senescence has emerged as a central hallmark of aging that contributes to age-related tissue dysfunction and the progression of age-related diseases (Ajoolabady et al., 2025). It was initially defined as a permanent state of cell-cycle arrest triggered by stressors such as telomere shortening, DNA damage and oxidative stress. Amongst other features, senescent cells exhibit morphological changes, persistent DNA damage, mitochondrial dysfunction and the secretion of pro-inflammatory molecules collectively termed the senescence-associated secretory phenotype (SASP). Senescence is increasingly recognized as an extremely dynamic and context-dependent cellular program characterized by substantial heterogeneity, with its features varying depending on the initial molecular trigger and the cell type (Cohn, Gasek, Kuchel, & Xu, 2023). While senescence has been extensively characterized in highly proliferative tissues, where repeated cell divisions lead to telomere attrition and replicative senescence, its characteristics and molecular and cellular consequences in the brain remain less well characterized.

Recent studies have shown that glial populations, such as microglia and astrocytes, can acquire shortened telomeres with age and exhibit a proinflammatory secretory profile and altered gene expression, driving chronic neuroinflammation and impaired glial support of neurons (Baker & Petersen, 2018). Importantly, even though neurons are largely post-mitotic, telomere dysfunction can still occur through DNA damage and inflammation, and studies have demonstrated that they can acquire a senescence-like phenotype (Jurk et al., 2012; Jurk et al., 2014). In fact, the relatively large metabolic demand of neurons coupled with their long lifespan and limited antioxidant defense seems to render them especially sensitive to DNA damage accumulation (Jurk et al., 2012). Additionally, senescence and mitochondrial dysfunction are tightly interconnected, with senescent cells displaying reduced respiratory capacity (Miwa, Kashyap, Chini, & von Zglinicki, 2022). Given the brain’s exceptionally high metabolic demand, such mitochondrial impairment can precipitate neuronal energy failure, with important consequences for neuronal activity and survival.

In our previous studies, we investigated how cellular senescence creates a permissive environment for AD pathology. Using cellular and mouse models, we established a causal link between telomere-induced senescence and intraneuronal Aβ accumulation (Suelves et al., 2023). Furthermore, we showed that telomere shortening accelerates tau pathology, including tau phosphorylation, truncation and aggregation, leading to exacerbated neuroinflammatory responses and neurodegeneration (Palomares et al., 2025). These findings support a central role for cellular senescence in linking aging-associated cellular stress to amyloid and tau pathologies, yet the underlying mechanisms remain incompletely understood. Moreover, although the removal of senescent cells using pharmacological treatments (senolytics) has shown benefits in multiple models (Baker & Petersen, 2018; Saez-Atienzar & Masliah, 2020), these treatments have not been effective when administered at advanced disease stages (Ng, Zhang, Li, & Baker, 2024), highlighting the importance of early intervention. Multi-omics approaches are key to capturing the complex interplay of transcriptional, proteomic, and cellular alterations contributing to pathological brain aging. Defining these molecular changes may reveal early mechanisms driving neurodegeneration and identify potential therapeutic targets for age-associated neurological disorders.

In the present study, we characterized the molecular landscape of brain senescence by conducting transcriptomic and proteomic analyses of brain tissue from Terc knockout mice. Pathway enrichment analysis highlighted a marked dysregulation of oxidative phosphorylation (OXPHOS)-related pathways. Functional analyses in mouse brain tissue further confirmed altered activity of mitochondrial respiratory chain complexes, with significant disturbances in brain energy homeostasis becoming apparent at later ages. In primary neurons derived from these mice, mitochondrial dysfunction was observed with increased ROS production, reduced ATP levels, and altered mitochondrial mass, although respiratory activity and neuronal viability remained largely preserved. In mouse brain tissue, our results highlighted a region-specific vulnerability of selected brain regions, where subtle alterations in neuronal survival were detected, suggesting that telomere attrition may prime specific neuronal populations for degeneration. Together, these findings identify mitochondrial dysfunction as an early and prominent consequence of telomere attrition in the brain, providing mechanistic insight into how telomere-induced senescence during pathological aging might lead to brain dysfunction and the onset of neurodegenerative conditions.

## 2 Results

### 2.1 Senescence markers accumulate in the brain of Terc knockout mice

Telomere attrition provides a robust in vivo model to study senescence-driven pathologies. Here, in agreement with our previous studies (Palomares et al., 2025; Suelves et al., 2023), we sought to validate whether the brains of Terc knockout mice display multiple established hallmarks of cellular senescence. As expected, telomere length measurements in cortical tissue revealed progressive telomere shortening across successive generations of Terc^-/-^ mice (G1-G3) at 5 months of age, indicating increased telomere dysfunction in later-generation animals (Figure 1A). Consistent with this finding, protein levels of γH2AX, a marker of DNA double-strand breaks (Rogakou, Pilch, Orr, Ivanova, & Bonner, 1998) and cellular senescence (Sedelnikova et al., 2004), progressively increased in hippocampal extracts from Terc^-/-^ mice compared to WT controls, reaching statistical significance in G2Terc^-/-^ and G3Terc^-/-^ animals (Figure 1B).

**Figure 1.**
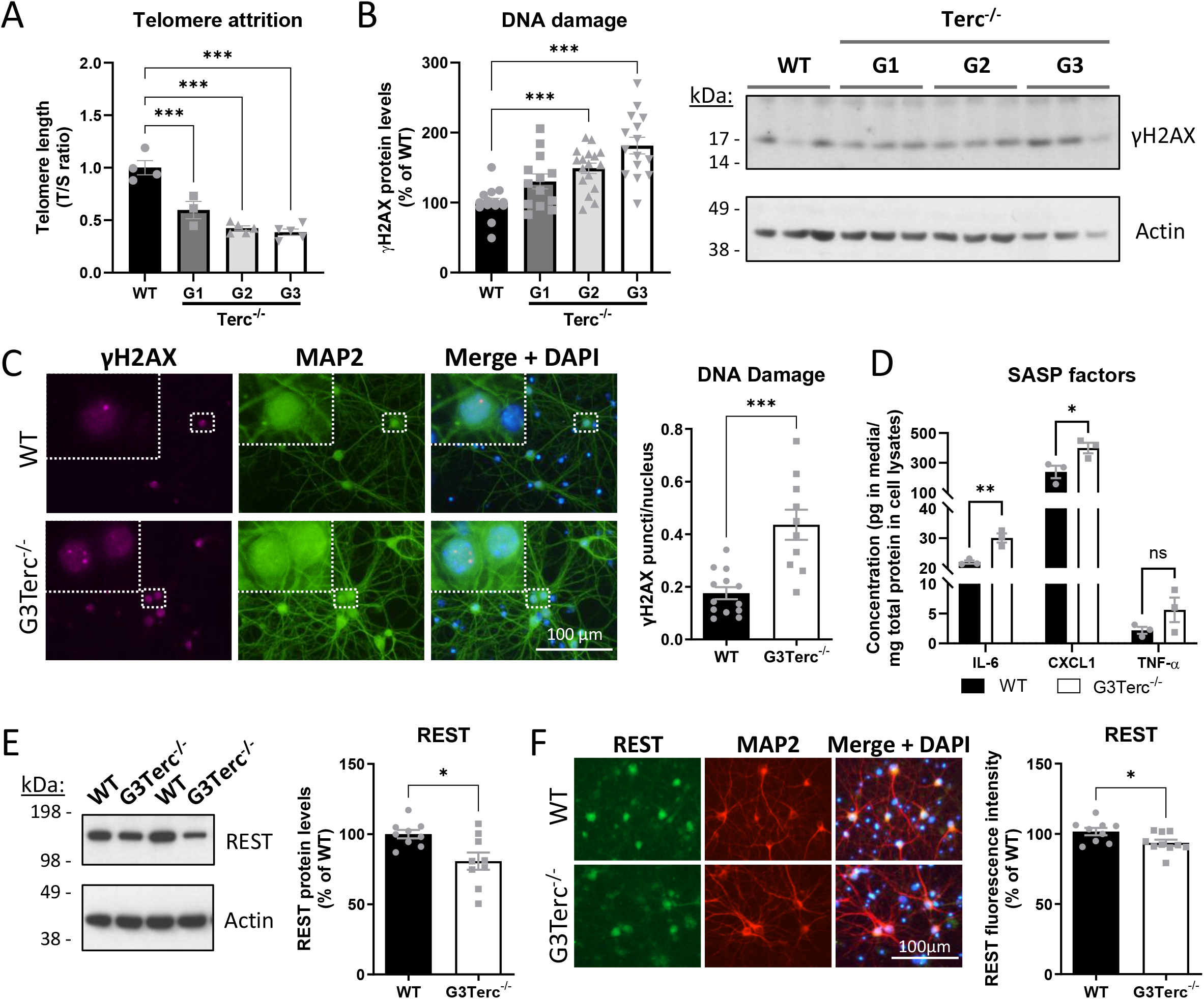
Cellular senescence markers are detected in the brains of telomerase-deficient mice. (A) qPCR analysis of relative telomere length in cortical samples from 5-month-old WT and successive generations (G1-G4) of Terc^-/-^ mice. Relative telomere length was calculated as the ratio (T/S) of telomere repeat copy number (T) to the copy number of a single-copy gene (S = 36B4). ***P < 0.001 (One-way ANOVA with Dunnett’s post-hoc analysis, n = 3-5 mice/group). (B) Western blot analysis of γH2AX levels in hippocampal samples from 5-month-old WT and successive generations (G1-G4) of Terc^-/-^ mice. Actin was used as a loading control, and levels in the WT group were normalized to 100%. ***P < 0.001 (One-way ANOVA with Dunnett’s post-hoc analysis, n = 14-17 mice/group). (C) Immunofluorescence analysis of the DNA damage marker γH2AX (purple) and the neuronal marker MAP2 (green) was performed in primary neurons derived from WT and G3Terc^-/-^ mice, cultured for 21 days in vitro (DIV21). The number of nuclear γH2AX foci per neuron was quantified. ***P < 0.001 (two-tailed Student’s t-test, n = 10-13 cultures/group). (D) Electrochemiluminescence immunoassay (ECLIA) quantification of IL-6, CXCL1, and TNF-α in conditioned media from DIV14 WT and G3Terc^-/-^ primary neurons. *P < 0.05, **P < 0.01 (two-tailed Student’s t-test, n = 3 cultures/group). (E) Western blot analysis of REST protein levels in WT and G3Terc^-/-^ primary neurons at DIV21. Actin was used as a loading control and levels in the WT group were normalized to 100%. *P < 0.05 (two-tailed Student’s t-test, n = 9 cultures/group). (F) Immunofluorescence analysis of REST (green) and the neuronal marker MAP2 (red) was performed in DIV21 primary neurons derived from WT and G3Terc^-/-^ mice. REST fluorescence intensity was assessed and levels in the WT group were set to 100%. *P < 0.05 (two-tailed Student’s t-test, n = 9-10 cultures/group). All data are presented as mean ± SEM.

Given the cellular heterogeneity of the brain and the ongoing debate as to whether terminally differentiated cells such as neurons can acquire senescent features (Baker & Petersen, 2018), we next assessed senescence-associated markers in primary neuronal cultures. Bright nuclear γH2AX foci were readily detected by immunofluorescence, and their number was significantly increased in G3Terc^-/-^ neurons (Figure 1C). Consistent with the acquisition of a SASP, conditioned medium from G3Terc^-/-^ neurons exhibited increased secretion of IL-6 and CXCL1, along with a trend toward elevated TNF-α levels, as determined by ECLIA analysis (Figure 1D). The Repressor Element-1 Silencing Transcription Factor (REST, also known as NRSF) is a transcriptional repressor that protects the aging brain against neurodegenerative stress and cognitive decline and is downregulated during pathological aging and AD (Lu et al., 2014). Consistent with previous observations (Piechota et al., 2016), REST protein levels progressively increased over time in WT neuronal cultures, as determined by Western blot and immunofluorescence analyses (Figure S1). In contrast, aged G3Terc^-/-^ primary neurons (DIV21) exhibited significantly reduced REST protein levels by both Western blot (Figure 1E) and immunofluorescence (Figure 1F), suggesting a compromised protective aging response.

Overall, these results confirm the presence of a senescent phenotype in brain cells of Terc^-/-^ mice, supporting the rationale for our subsequent analyses.

### 2.2 Multi-omic analyses of Terc knockout mice identify mitochondrial dysfunction as a central hallmark of brain senescence

Understanding how cellular senescence contributes to brain aging and neurodegeneration has become a major research priority. However, the identification of senescent brain cells remains challenging due to the heterogeneity of senescence phenotypes and the lack of universal brain-specific senescence markers (Baker & Petersen, 2018). To address this, we used Terc knockout mice as a model of telomere-driven senescence and performed comprehensive multi-omic analyses to identify robust molecular signatures associated with brain aging.

To characterize the transcriptomic profile associated with telomere-induced senescence, we performed RNA-sequencing (RNA-seq) analyses on hippocampal tissue from 5-month-old WT and G3Terc^-/-^ mice (4 mice/genotype). Principal component analysis (PCA) revealed a clear segregation of the samples by genotype (Figure 2A). Consistent with this separation, differential expression analysis identified 244 differentially expressed genes (DEGs) between WT and G3Terc^-/-^ mice, comprising 127 upregulated and 117 downregulated genes (Figure 2B and Table S1, DEGs ranked by adjusted p-value). Differences in mRNA expression levels of a subset of DEGs were validated by RT-qPCR (Figure S2). Specifically, upregulation of *Cdkn1a* (p21) and *Ccnd1* (Cyclin D1) expression reinforces the presence of cell cycle-associated alterations in our model, while downregulation of *Syt1* (Synaptotagmin 1), a Ca²⁺-sensing protein essential for synaptic vesicle exocytosis and neurotransmitter release, might contribute to synaptic dysfunction. Importantly, reduced SYT1 levels have been reported in multiple brain regions of AD patients (Öhrfelt et al., 2016), supporting its relevance as an early marker of synaptic dysfunction in age-associated neurodegeneration.

**Figure 2.**
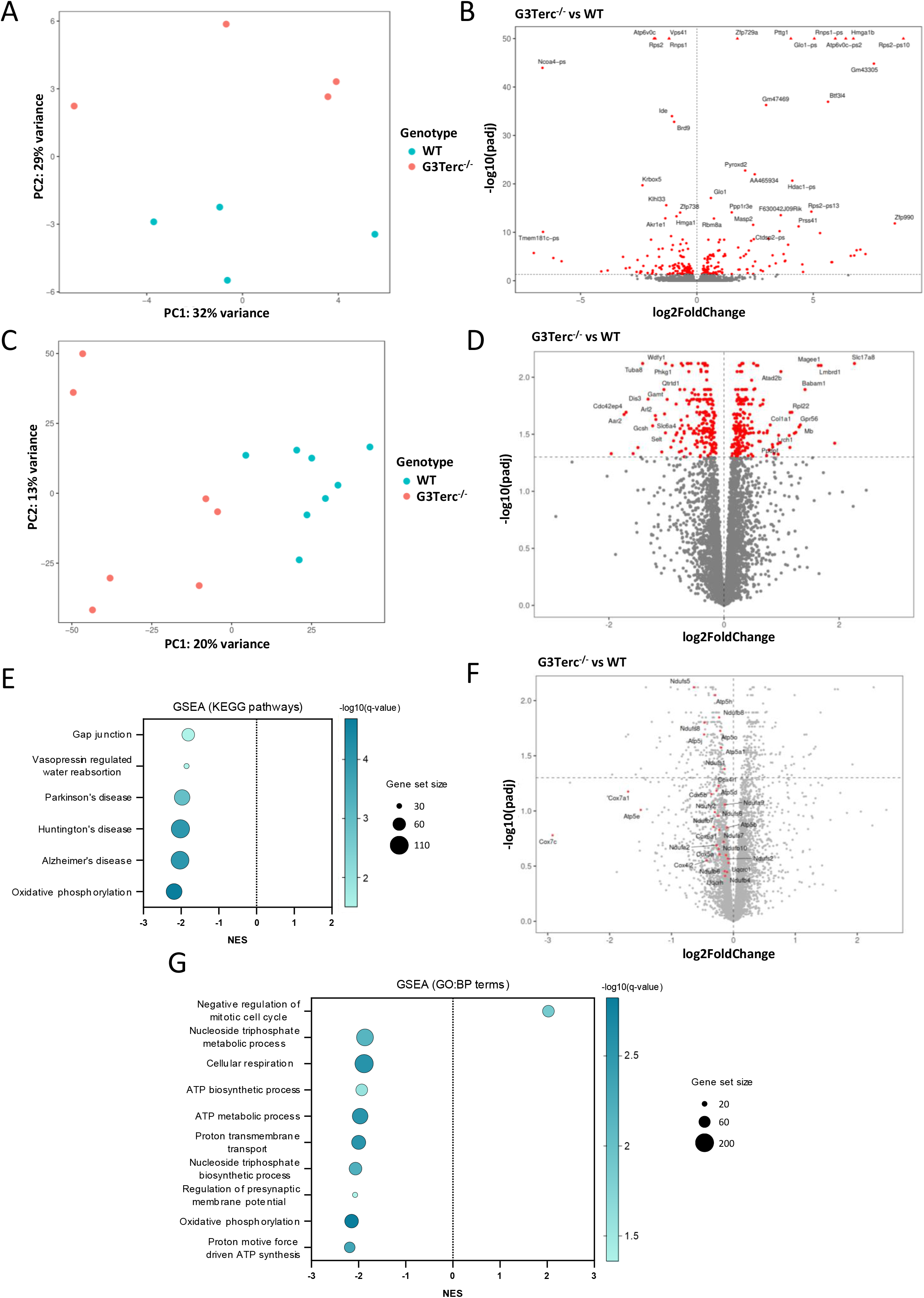
Transcriptomic and proteomic profiling of brain senescence in telomerase-deficient mice. RNA sequencing (RNAseq) (A-B) and LC-MS/MS (C-G) were performed on hippocampal tissue from 5-month-old WT and G3Terc^-/-^ mice to characterize molecular alterations associated with telomere-induced senescence. (A, C) Principal component analysis (PCA) of transcriptomic and proteomic datasets, respectively. (B, D) Volcano plots showing differential gene and protein expression in G3Terc^-/-^ hippocampi relative to WT controls. Differentially expressed genes (DEGs) and differentially expressed proteins (DEPs) were identified using a false discovery rate (FDR)-adjusted p-value < 0.05. In B, data points for genes with p-adjusted values < 10^-50^ are shown as triangles at −log10(padj) = 50. In D, DEPs are represented by their corresponding gene symbols. (E, G) Gene set enrichment analysis (GSEA) of LC-MS/MS data was performed using Kyoto Encyclopedia of Genes and Genomes (KEGG) pathways (E) or Gene Ontology Biological Process (GO:BP) terms (G). Bubble plots depict enriched pathways and their normalized enrichment score (NES), with bubble size indicating the number of genes per pathway (set size) and color representing statistical significance (adjusted p-value). The top ten enriched GO:BP terms are shown. (F) Volcano plot highlighting (in red) the 31 leading-edge genes shared among the four most significant KEGG pathways identified by GSEA: Oxidative phosphorylation, Alzheimer’s disease, Parkinson’s disease, and Huntington’s disease.

To gain insight into the biological pathways altered in G3Terc^-/-^ brains, we performed Gene Set Enrichment Analysis (GSEA) using GO (Gene Ontology) and KEGG (Kyoto Encyclopedia of Genes and Genomes) databases. However, no pathways reached statistical significance after correcting for multiple testing. These findings suggest that Terc knockout brains exhibit relatively mild transcriptional alterations.

To further explore senescence signatures at the protein level, we performed Liquid Chromatography-Tandem Mass Spectrometry (LC-MS/MS)-based proteomic analyses on hippocampal extracts from an expanded sample set of WT and G3Terc^-/-^ mice at 5 months of age (7-8 mice/genotype). This cohort included the animals previously analyzed by RNA-seq, plus additional animals from each genotype. PCA representation revealed again a clear segregation of samples by genotype (Figure 2C). A total of 433 differentially expressed proteins (DEPs) were detected between G3Terc^-/-^ and WT brains, including 219 up-regulated and 214 down-regulated ones (Figure 2D and Table S2). Of these, only 28 DEPs exhibited an absolute log2 fold change greater than 1 (|log2FC| > 1), indicating that the majority of protein expression changes were of modest magnitude. Despite this, GSEA pathway analysis using the KEGG database identified Oxidative phosphorylation (OXPHOS) as the most strongly negatively enriched and statistically significant pathway, followed by pathways associated with neurodegenerative disorders, including Alzheimer’s disease, Huntington’s disease, and Parkinson’s disease (Figure 2E and Table S3). These pathways shared a substantial proportion of their core enrichment genes, with 31 genes being common to all four pathways (Figure 2F). The majority of these genes encode key components of the mitochondrial respiratory chain, spanning ETC Complexes I, III, IV, and V. This redundancy between KEGG gene sets suggests that the observed enrichment is driven by mitochondrial dysfunction, with the neurodegenerative disease pathways representing alternative annotations of the same underlying transcriptional program. Additionally, the KEGG pathways Vasopressin regulated water reabsorption and Gap junction appeared downregulated, although with lower statistical significance and weaker enrichment scores. Closer inspection of their core-enrichment genes suggested alterations in cytoskeletal organization and intracellular signaling in Terc knockout brains.

GSEA analysis using Gene Ontology Biological Process (GO:BP) terms reinforced these findings, revealing a significant downregulation of processes associated with mitochondrial bioenergetics, including OXPHOS, aerobic/cellular respiration, electron transport chain and ATP production, largely driven by the same genes that appeared in the KEGG analysis (Figure 2G and Table S3). Additionally, Regulation of presynaptic membrane potential was significantly downregulated, driven by reduced expression of excitatory glutamatergic receptors (e.g., *Grik2, Gria3*), inhibitory GABA receptors (e.g., *Gabbr1, Gabra5*), and neuronal ion channels (e.g., *Kcnj9*, *Kcnj3, Scn1a, Kcnc2*), suggesting alterations in neuronal communication in G3Terc^-/-^ hippocampi. Finally, GSEA using GO:BP terms identified the term Negative regulation of mitotic cell cycle as positively enriched, involving regulators of the DNA damage response (e.g., *Babam1*, *Rad50, Ppp1r10*), cell cycle progression (e.g., *Tfdp1*, *Brinp2*, *Cdkn1b*), and ubiquitin-mediated protein turnover (e.g., *Nae1, Fbxo7*, *Psmg2*), among others, consistent with altered cell cycle progression and a senescence-like phenotype in G3Terc^-/-^ mice.

Given the strong mitochondrial signature observed in the proteomic analysis, we next performed a targeted analysis of the OXPHOS pathway in the RNA-seq dataset to determine whether these OXPHOS-associated alterations were also evident at the transcriptional level. Although OXPHOS did not reach statistical significance after multiple-testing correction in the initial unbiased GSEA (data not shown), targeted analysis of this pathway revealed a significant and negative enrichment signal (NES = -1.71, p-value = 0.0069). Comparison of the OXPHOS core-enrichment genes identified in the proteomic and RNA-seq analyses revealed eight genes shared between the two datasets (*Cox15*, *Cox5a*, *Cox5b*, *Cox7c*, *Ndufa9*, *Ndufb6*, *Ndufb7*, and *Ndufb10*), all encoding components of mitochondrial respiratory chain Complexes I or IV. Together, these findings suggest that the observed OXPHOS alterations in Terc knockout brains are more pronounced at the protein level and exhibit relatively limited overlap between transcriptional and proteomic changes.

Consequently, to further assess the consistency between the transcriptomic and proteomic alterations, differential expression statistics from both datasets were compared. Strikingly, RNA and protein changes showed little correlation (Pearson’s r = 0.06), with only three genes significantly and concordantly dysregulated at both levels: *Rbm8a*, *Wdfy1* and *Ggps1* (Figure S3).

Together, these findings indicate a predominant impairment of mitochondrial bioenergetic function and cellular energy production in Terc knockout brains, particularly at the protein level.

### 2.3 Functional characterization reveals impaired OXPHOS activity and energy depletion in Terc-deficient brains

Given that multi-omics analyses revealed a significant downregulation of mitochondrial function and the OXPHOS pathway in senescent brains, we sought to validate these findings by assessing mitochondrial electron transport chain (ETC) activity. OXPHOS occurs at the inner mitochondrial membrane, where five multi-subunit protein complexes (I-V) and two electron carriers cooperate to generate ATP. To evaluate the activity of the mitochondrial complexes, functional mitochondria were freshly isolated from adult mouse brains and analyzed using the electron flow assay on the Seahorse XF96 platform (Agilent). This assay measures oxygen consumption rates (OCR) following the sequential addition of specific substrates and inhibitors targeting individual ETC complexes. We first analyzed the activity of mitochondria isolated from the hippocampus and cortex of 5-month-old WT and G3Terc^-/-^ mice (Figure 3A). In hippocampal mitochondria, the activities of complexes II, III, and IV were significantly reduced in G3Terc^-/-^ mice, while complex I activity showed a similar downward trend. In cortical mitochondria, complex II activity was significantly reduced, whereas complexes I, III, and IV also displayed decreased activity trends.

**Figure 3.**
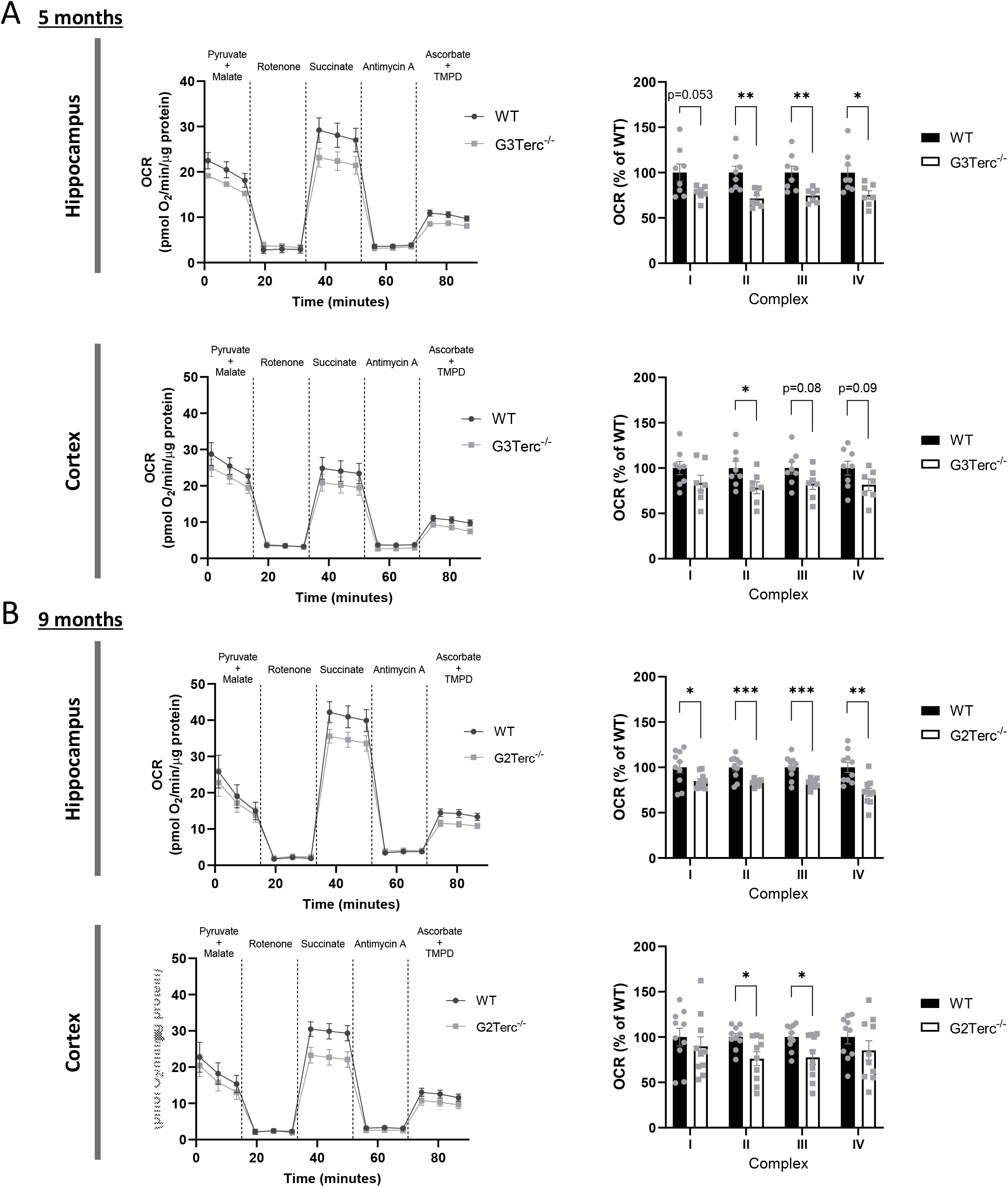
Altered activity of the mitochondrial ETC complexes during brain senescence. (A, B) Electron flow assay in mitochondria isolated from hippocampal and cortical extracts of 5-month-old WT and G3Terc^-/-^ mice (A) and 9- month-old WT and G2Terc^-/-^ mice (B). Curves of oxygen consumption rates (OCR) are shown. Mitochondrial complex activities were calculated from changes in OCR following sequential injections of complex substrates and inhibitors: Complex I activity was determined by subtracting the OCR after rotenone injection from the OCR measured after pyruvate and malate incubation. Complex II activity was calculated by subtracting the OCR after rotenone injection from the OCR after succinate addition. Complex III activity was determined by subtracting the OCR after antimycin A injection from the OCR after succinate addition. Finally, complex IV activity was calculated by subtracting the OCR after antimycin A injection from the OCR measured following ascorbate and TMPD addition. *P < 0.05, **P < 0.01, ***P < 0.001 (two-tailed Student’s t-test, n = 7-10 mice/group). All data are presented as mean ± SEM.

Senescence is a dynamic process whose effects may change over time as the organism’s capacity to adapt to pathological alterations declines with age. To better characterize the impact of brain senescence in an aged context, later disease stages needed to be evaluated. However, G3Terc^-/-^ mice rarely survived beyond 5 months of age. To overcome this limitation, we turned to the G2Terc^-/-^ model, which has a longer lifespan while still exhibiting significant telomere shortening, classical senescent features and enhanced chronic inflammation, as previously demonstrated by our group (Palomares et al., 2025). We therefore analyzed G2Terc^-/-^ mice at 9 months of age. Similarly to what was observed in 5-month-old mice, aged senescent brains exhibited a comparable reduction in the activity of complexes I-IV in the hippocampus, as well as complexes II and III in the cortex (Figure 3B). Overall, our findings indicate impaired ETC complex activity in young and aged Terc^-/-^ mouse brains.

To determine the mechanisms underlying this functional impairment, we first assessed whether it could be explained by defects in mitochondrial complex assembly. However, the absence of major changes in representative OXPHOS subunit levels in mitochondrial protein extracts from these mice (Figure S4) argues against a substantial loss or destabilization of the respiratory chain complexes. Moreover, although GSEA analysis of proteomic data identified the OXPHOS pathway as negatively enriched (Figure 2E-G), fold changes were very mild, suggesting that consistent but mild downregulation of many OXPHOS subunits underlies these effects, rather than a loss of any individual subunit. We also considered whether mitochondrial genome integrity itself might be compromised. Using a polymerase-based assay, in which equal amounts of input DNA are amplified across samples and damage is inferred from the relative reduction in the long mtDNA amplification product, we detected significant mitochondrial DNA (mtDNA) damage in G3Terc^-/-^ mice (Figure S5A). In parallel, mtDNA content was evaluated by qPCR quantification of mitochondrially-encoded genes, revealing no differences and therefore providing a normalization control for the mtDNA damage results (Figure S5B). Given this mtDNA damage, we next assessed whether this was associated with transcript-level alterations in the 13 mitochondrial-encoded genes using our RNA-seq data. Although no individual gene reached significance on its own (Table S1), the mitochondrial-encoded gene set revealed a significant negative enrichment (NES = -2.23, p value = 7.4 × 10^-6^), suggesting a subtle but coordinated transcriptional downregulation of the mitochondrial genome that could be driven by this damage (Figure S6). Overall, our findings suggest that the reduced respiratory capacity observed in telomere-dysfunctional brains may stem from impaired mitochondrial genome integrity, accompanied by mild but consistent protein-level downregulation of OXPHOS components, rather than changes in mitochondrial abundance.

The observed mitochondrial alterations could translate into changes in the energetic state of the brain. Neurons are highly energy-demanding cells that rely heavily on mitochondrial ATP production to sustain physiological functions. During cellular metabolism, ATP is progressively converted into ADP and AMP, making the abundance of these nucleotides a reliable indicator of cellular bioenergetic state. To evaluate energy metabolism in our senescence models, ATP, ADP, and AMP levels were quantified by High-Performance Liquid Chromatography (HPLC). Measurements were first obtained in hippocampal and cortical tissues from 5-month-old WT and G3Terc^-/-^ mice. Surprisingly, despite the reduced ETC activity observed in these animals, neither brain region displayed significant differences in ATP, ADP, or AMP levels, nor in the calculated energy charge, compared to WT mice (Figure 4A). These results may suggest the presence of compensatory mechanisms in young G3Terc^-/-^ mice that preserve global brain energy homeostasis despite reduced mitochondrial respiratory activity.

**Figure 4.**
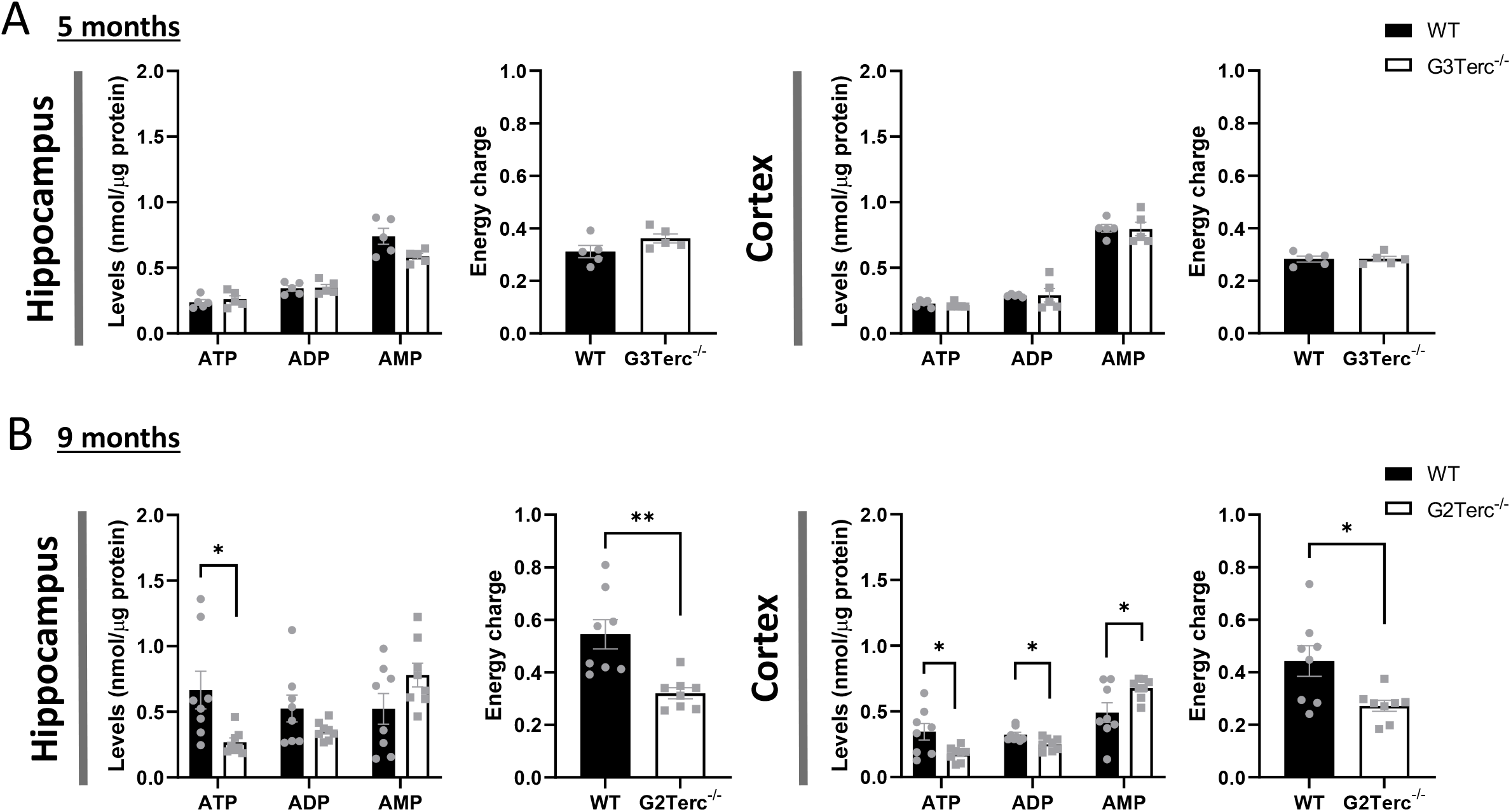
Altered brain energy homeostasis during brain senescence at advanced ages. (A,B) HPLC analysis of ATP, ADP, and AMP levels in the hippocampus and cortex of 5-month-old WT and G3Terc^-/-^ mice (A) and 9-month-old WT and G2Terc^-/-^ mice (B). Nucleotide levels are expressed per μg of tissue protein extract, and the energy charge (EC) was calculated as EC = ([ATP] + 0.5[ADP]) / ([ATP] + [ADP] + [AMP]). *P < 0.05, **P < 0.01 (Two-tailed Student’s t-test, n = 5-8 mice/group). All data are presented as mean ± SEM.

To determine whether aging exacerbates energetic dysfunction in the context of senescence, HPLC measurements were performed in hippocampal and cortical tissue from 9-month-old WT and G2Terc^-/-^ mice (Figure 4B). In contrast to the previous findings, ATP levels were significantly reduced in both the hippocampus and cortex of G2Terc^-/-^ mice, while the cortex displayed an additional reduction in ADP accompanied by a significant increase in AMP levels. Consistent with these alterations, overall energy charges were significantly depleted in both hippocampal and cortical regions compared to WT controls. Overall, these findings support a model in which reduced mitochondrial OXPHOS activity represents an early feature of telomere-induced senescence, while its consequences for cellular bioenergetics become progressively apparent with aging.

### 2.4 Accumulation of dysfunctional mitochondria in primary neurons from Terc knockout mice

As measurements of OXPHOS activity and adenine nucleotide levels were performed on whole-tissue extracts, we next sought to better characterize the cell-type-specific alterations underlying these bioenergetic defects by focusing on primary neuronal cultures derived from G3Terc^-/-^ mice. We previously validated that these neurons exhibit several classical hallmarks of cellular senescence (Figure 1 and (Suelves et al., 2023)). In contrast to brain tissue observations (Figure S5), senescent primary neurons at DIV14 showed a marked increase in mitochondrial mass, as evidenced by both elevated mtDNA content (Figure 5A) and by increased TOM20 protein levels (Figure 5B). Interestingly, TOM20 expression remained unchanged in primary astrocytes derived from G3Terc^-/-^ mice, indicating that mitochondrial accumulation is a neuron-specific feature of the senescent phenotype (Figure 5B). To further characterize neuron-specific bioenergetic alterations, reactive oxygen species (ROS) production was assessed using the CM-H2DCFDA fluorescent probe as an indirect indicator of oxidative stress and mitochondrial dysfunction. Senescent neurons at DIV14 exhibited significantly elevated ROS levels, as demonstrated both by fluorescence microscopy (Figure 5C) and microplate-based quantification (Figure 5D). In parallel, ATP levels measured using the CellTiter-Glo luminescent assay were significantly reduced in DIV14 G3Terc^-/-^ neurons compared to WT controls (Figure 5E), indicating the presence of a bioenergetic deficit. Mitochondrial respiratory function was subsequently evaluated in DIV14 primary neurons derived from WT and G3Terc^-/-^ mice using the Mito Stress Test assay on the Seahorse XF platform. Surprisingly, no significant differences in OCR-derived parameters were detected between genotypes (Figure 5F), including basal respiration, ATP production, maximal respiration, and spare respiratory capacity. Consistent with this apparent preservation of mitochondrial respiration, cell viability assessed by an MTS-based assay revealed no significant differences between genotypes (Figure 5G). To evaluate neuronal activity, primary neurons were stimulated with bicuculline (50 μM) or vehicle solution for 2 h, followed by Western blot analysis of c-Fos, Egr1, and NPAS4 protein levels (Figure S7). These activity-dependent immediate early-gene (IEG) products are rapidly induced after neuronal activation and are widely used as markers of transcriptional responses to synaptic activation (Yap & Greenberg, 2018). Bicuculline treatment strongly induced the protein expression of all three IEGs in both WT and G3Terc^-/-^ neurons, with Two-way ANOVA analysis detecting a significant main effect of treatment. However, no significant main effect of genotype or treatment x genotype interaction was detected, suggesting that activity-dependent responses are preserved in G3Terc^-/-^ neurons.

**Figure 5.**
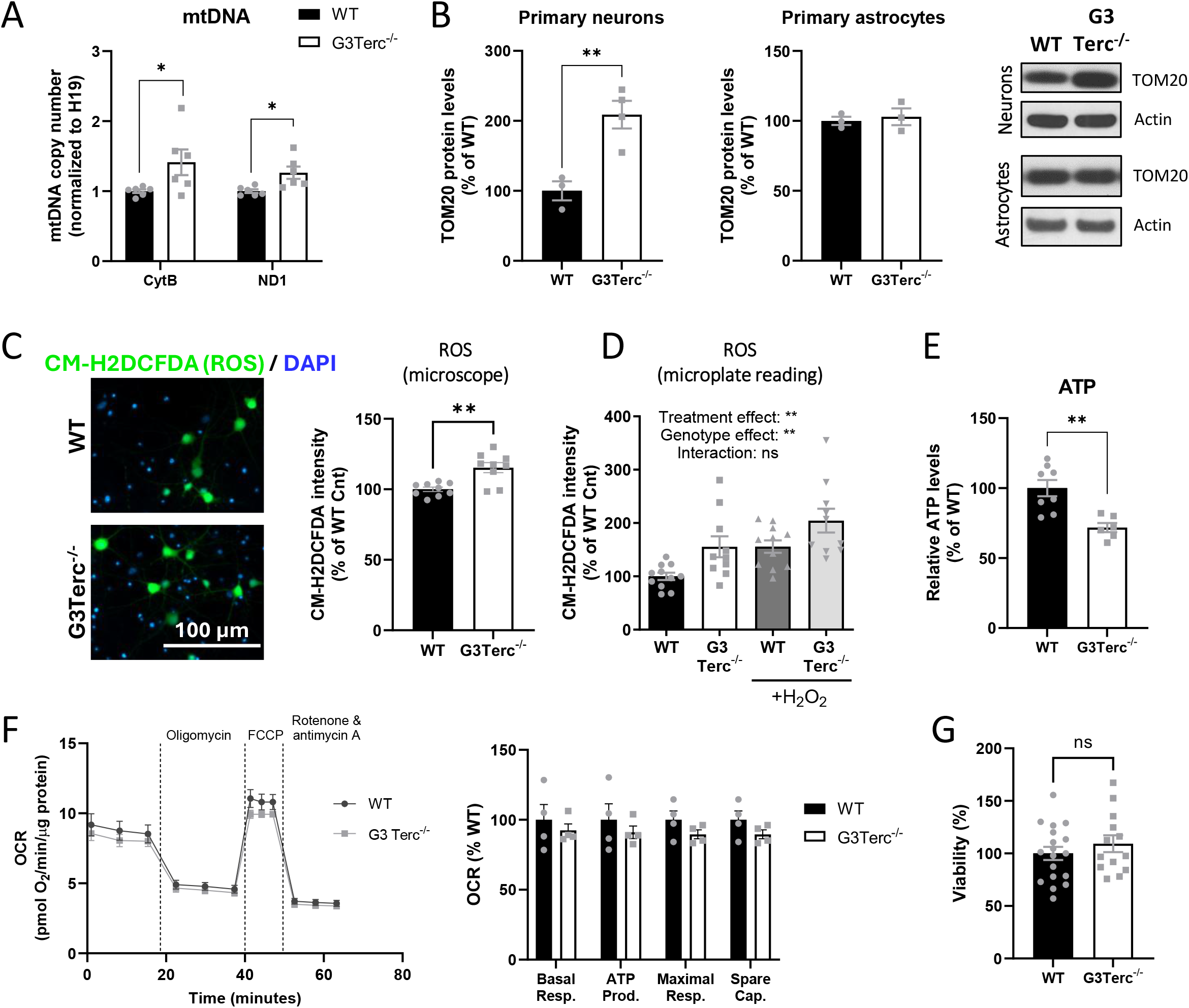
Senescent primary neurons accumulate dysfunctional mitochondria despite preserved viability. (A) qPCR analysis of mitochondrial DNA (mtDNA) content (*CytB* and *Nd1* genes) in DIV14 primary neurons derived from WT and G3Terc^-/-^ mice. *P < 0.05 (Two-tailed Student’s t-test, n = 6 cultures/group). (B) Western blot analysis of TOM20 protein levels in DIV14 primary neurons (left) or DIV23 primary astrocytes (right) derived from WT and G3Terc^-/-^ mice. **P < 0.01 (Two-tailed Student’s t-test, n = 3-4 cultures/group). (C, D) Production of reactive oxidative species (ROS) in DIV14 WT and G3Terc^-/-^ primary neurons was assessed by measuring CM-H2DCFDA fluorescence intensity using either fluorescence microscopy (C) or a plate reader (D). For plate reader measurements, a subset of neurons was treated with 1 mM hydrogen peroxide (H₂O₂), a well-established inducer of ROS production, as a positive control. For C,**P < 0.01 (Two-tailed Student’s t-test, n = 9 cultures/group). For D, **P < 0.01 (Two-way ANOVA with Tukey’s post-hoc analysis, n = 10-11 cultures/group). (E) Relative ATP levels in DIV14 WT and G3Terc^-/-^ primary neurons were measured using a luminescent assay (CellTiter-Glo). **P < 0.01 (Two-tailed Student’s t-test, n = 6-8 cultures/group). (F) Mitochondrial respiration was evaluated in WT and G3Terc^-/-^ primary neurons at DIV14 with the Seahorse XFe96 Analyzer and the Mito Stress kit. The oxygen consumption rate (OCR) profile is shown, with vertical dashed lines indicating the sequential addition of oligomycin (2 μM; CV inhibitor), FCCP (0.5 μM; mitochondrial membrane potential uncoupler), and rotenone/antimycin A (0.5 μM each; complex I and complex III inhibitors, respectively). Basal respiration, ATP-linked respiration, maximal respiration, and spare respiratory capacity were calculated according to the manufacturer’s recommended protocol. ns (two-tailed Student’s t-test, n = 4 cultures/group). (G) An MTS-based assay was used to assess viability on DIV14 WT and G3Terc^-/-^ primary neurons. ns (two-tailed Student’s t-test, n = 13-18 cultures/group). All data are presented as mean ± SEM.

Altogether, these findings suggest that senescent neurons accumulate dysfunctional mitochondria characterized by increased mitochondrial mass, elevated oxidative stress, and reduced ATP levels. The preservation of global respiratory activity and activity-dependent transcriptional responses despite these alterations may reflect a compensatory increase in mitochondrial content aimed at sustaining cellular energy demands in the context of mitochondrial dysfunction.

### 2.5 Selective neuronal loss in vulnerable brain regions of Terc knockout mice

As impaired mitochondrial activity and energetic imbalance can ultimately compromise neuronal function and integrity, we next investigated whether senescent brains exhibited alterations in neuronal survival and tissue organization. To this end, neuronal density and overall brain morphology were assessed by immunofluorescent staining using an anti-NeuN antibody, a marker of mature neurons (Figure 6A). NeuN-positive nuclei were automatically quantified within anatomically defined regions of the cortex and hippocampus using the QuPath software (Figure 6B). Overall, no signs of gross brain atrophy were detected. However, detailed regional analyses revealed subtle but significant reductions in neuronal abundance in specific vulnerable areas. While the CA1 and CA3 hippocampal subregions were largely preserved, the subiculum displayed a decrease in NeuN-positive cell counts compared to WT mice. Similarly, a reduction in neuronal numbers was observed in cortical layer V. These findings are consistent with previous observations from our group (Palomares et al., 2025; Suelves et al., 2023), reinforcing the notion that telomere-driven senescence leads to selective neuronal vulnerability rather than widespread neurodegeneration in the absence of additional pathological stressors.

**Figure 6.**
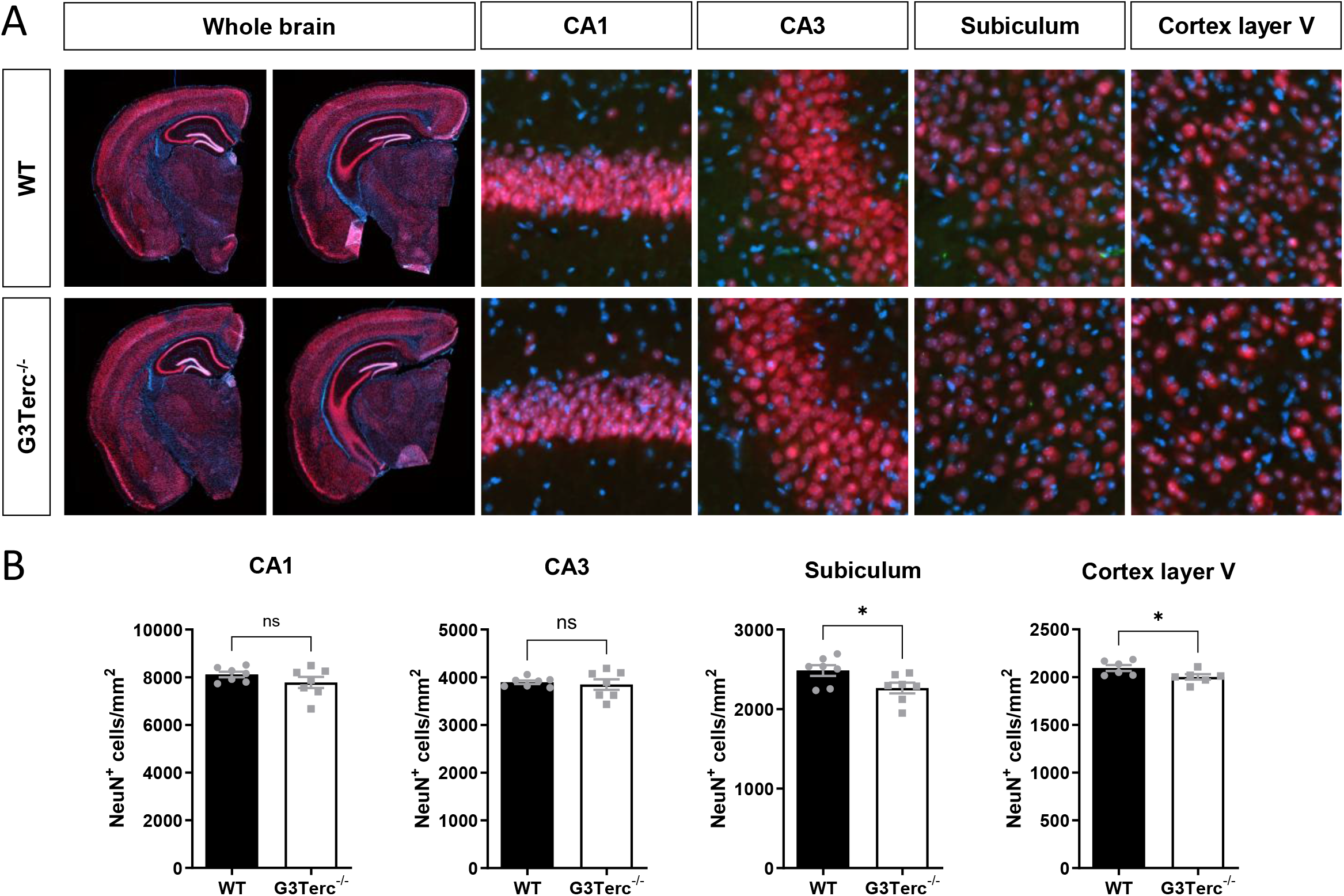
Telomere shortening mildly impairs neuronal viability in adult mice in a region-specific manner. (A) Representative Immunofluorescence images of the neuronal marker NeuN (red) in 5/6-month-old WT and G3Terc^-/-^ mice. (B) Quantification of NeuN-positive cell density in the CA1, CA3, subiculum, and layer V of the cerebral cortex shown in A.* P < 0.05 (two-tailed Student’s t-test, n = 7 mice/group). All data are presented as mean ± SEM.

## 3 Discussion

Increasing evidence from our group and others indicates that the accumulation of senescent cells during aging contributes to the onset and progression of neurodegenerative pathologies (Baker & Petersen, 2018; Palomares et al., 2025; Saez-Atienzar & Masliah, 2020; Suelves et al., 2023), but the underlying mechanisms remain poorly understood. Our study provides a comprehensive landscape of the transcriptomic and proteomic alterations induced by telomere-driven senescence in the mouse brain, identifying mitochondrial pathways as among the most prominently affected processes and proposing potential novel markers of senescence. This molecular signature was validated by functional analyses, which revealed ETC complex activity and reduced energetic status in brain tissue. In vitro experiments in primary neurons further confirmed the presence of mitochondrial dysfunction and impaired energy metabolism through cell-autonomous mechanisms. Finally, in vivo analyses indicated that these alterations are associated with modest, region-specific neuronal loss.

Telomere shortening is an evolutionarily conserved aging mechanism that has been extensively characterized in multiple species. It triggers a persistent DNA damage response (DDR), which constitutes one of the major drivers of cellular senescence (d’Adda di Fagagna et al., 2003). Thus, telomerase-deficient mouse models, including Terc knockout mice, are well-established systems for studying cellular senescence in vivo (Rossiello, Jurk, Passos, & d’Adda di Fagagna, 2022; Rudolph et al., 1999). However, whether physiological brain aging is accompanied by progressive telomere shortening remains debated, particularly because mature neurons are post-mitotic and no longer undergo DNA replication (Schreglmann et al., 2023; Tomita et al., 2018). Nevertheless, telomere dysfunction capable of inducing senescence can arise independently of telomere length; telomeric DNA is intrinsically more susceptible to oxidative damage due to its guanine-rich repetitive sequences, and DNA repair is less efficient at telomeres than in the rest of the genome (Rossiello et al., 2022). Additionally, chronic inflammation has been shown to induce telomere dysfunction and accelerate aging in mice, even in the absence of extensive telomere shortening (Jurk et al., 2014). Thus, our findings in Terc knockout mouse brains provide a framework to evaluate the consequences of senescence independently of whether telomere attrition is a primary driver of brain aging. Moreover, recent evidence suggests that telomere attrition may be exacerbated by AD pathology itself. In particular, shorter telomere length was observed in brain tissue from the APP/PS1 amyloid mouse model (X. Guo et al., 2024). In this context, we propose that an increasing pathological burden would further accelerate telomere erosion, which would in turn downregulate mitochondrial activity and establish a feed-forward mechanism that amplifies neuronal dysfunction during disease progression.

Understanding the mechanisms that drive age-associated pathological changes is essential for identifying pathways that may be targeted therapeutically, but progress has been hindered by the heterogeneity of aging and the diversity of experimental models. Cell type-specific changes add further complexity, often making it difficult to generalize classical aging markers that have been predominantly characterized in proliferative cells. An ongoing debate in the field is whether post-mitotic cells such as neurons can exhibit classical hallmarks of cellular senescence. Our findings in primary neuronal cultures suggest that neurons can develop several senescence-associated hallmarks, including increased SASP production (IL-6 and CXCL1) and enhanced DNA damage accumulation, visualized by elevated γH2AX nuclear foci. Previous data from our group also demonstrated increased expression of cell cycle regulatory genes (Suelves et al., 2023). These findings are in agreement with previous studies (Jurk et al., 2012), showing increased IL-6 and γH2AX levels in aged neurons and Terc^-/-^ brains. Moreover, our data indicate that aged (21DIV) G3Terc^-/-^ neurons exhibit reduced protein levels of REST, a more recently recognized marker of neuronal aging (Lu et al., 2014; Piechota et al., 2016). REST is highly expressed during embryonic development, where it represses neuronal gene expression in non-neuronal cells and neural progenitors (Johnson et al., 2008). Its expression declines as neurons differentiate and mature but is re-induced in the aging brain, where it acquires neuroprotective functions. Notably, loss of REST expression is associated with cognitive deficits and AD (Lu et al., 2014). Thus, the decreased REST expression observed in our telomere-deficient neurons provides further evidence that telomere attrition induces molecular changes associated with pathological brain aging.

Recent advances in single-cell omics have enabled cell-type-specific resolution of complex biological processes, including brain aging (Sun, Nagvekar, Pogson, & Brunet, 2025). Due to the fragility of brain cells and the difficulty of isolating intact cells, single-nucleus approaches are often used, but they are biased toward nuclear and nascent transcripts and therefore underrepresent cytoplasmic RNAs, including mitochondrially encoded genes. Our bulk omics analyses provide an unbiased assessment of transcriptomic and proteomic alterations, avoiding potential artifacts introduced by cell or nuclear isolation. We identify mitochondrial dysfunction, characterized by reduced OXPHOS function and energy hypometabolism, as a major process affected by telomere-driven brain senescence, particularly at the protein level. Consistent with our findings, defects in OXPHOS function have been reported in multiple models of senescence (Miwa et al., 2022), including decreased activity of ETC complexes in isolated heart and liver mitochondria from G2Terc^-/-^ mice (Sahin et al., 2011), although most evidence derives from non-neuronal cells. Importantly, similar defects in OXPHOS and energy metabolism are consistently observed as early and prominent features in AD patients (W. Wang, Zhao, Ma, Perry, & Zhu, 2020), suggesting that mitochondrial dysfunction arising during pathological aging (with induction of senescence processes) may act as an early driver of disease pathogenesis. In accordance with this, KEGG GSEA analysis revealed enrichment of neurodegenerative disease pathways, including AD, largely driven by overlapping core enrichment genes associated with the mitochondrial respiratory chain. Future studies should validate these alterations, particularly in Nduf, Cox, and Atp5 family members, and determine their contribution to the observed functional impairments.

Beyond mitochondrial alterations, proteomic analyses also revealed positive enrichment of the gene set corresponding to Negative regulation of mitotic cell cycle, supporting the activation of a replicative senescence program in the brain. A gene set associated with presynaptic membrane potential was also found to be downregulated, mainly driven by reduced expression of excitatory glutamatergic receptors, inhibitory GABA receptors, and neuronal ion channels, suggesting a simultaneous impairment of excitatory and inhibitory synaptic signaling during pathological brain aging. Future studies should assess whether these molecular alterations translate into neuronal activity deficits using electrophysiology or calcium imaging.

The omics data indicate that brain senescence is characterized by numerous but subtle transcriptomic and even more abundant proteomic alterations. Importantly, such alterations correlate very weakly with each other, which may indicate the contribution of post-transcriptional or post-translational dysregulation. Consistently, senescent human cells in vitro showed robust SASP activation at the transcript level, whereas the corresponding secretome exhibited limited concordance with mRNA abundance (Sullivan et al., 2021). Additionally, the correlation between mRNA and protein levels has been shown to decline with age in the brain of humans and rhesus macaques, an effect enriched among genes involved in mitochondrial function and likely caused by RNA-binding proteins (Wei et al., 2015). Another study found that, despite an overall slowing of protein turnover in the aged mouse brain, a subset of mitochondrial proteins appears to be shorter-lived (Kluever et al., 2022). Deciphering whether similar mechanisms underlie the transcript-protein discordance observed here in the context of telomere-driven senescence represents an important avenue for future investigation.

In telomerase-deficient mice, although changes in individual OXPHOS components were modest and no global structural defects of the mitochondrial respiratory chain were detected, electron flow data revealed an approximately 20% reduction in the activity of multiple ETC complexes. These findings suggest that minor changes in the abundance of multiple respiratory chain components are sufficient to impair mitochondrial respiration. In addition, OXPHOS activity is known to be regulated by multiple post-translational mechanisms, including post-translational modifications and the assembly of mitochondrial supercomplexes (Hofer & Wenz, 2014; Lopez-Fabuel et al., 2016), which warrant further investigation. Even a relatively small reduction (20%) in mitochondrial function may have important consequences in the brain, which accounts for only ∼2% of total body mass yet consumes approximately 20% of the body’s oxygen (Raichle & Gusnard, 2002). In our model, telomere-induced senescence produced only mild neuronal loss in selected cortical and hippocampal populations, indicating that mitochondrial dysfunction initially manifests as functional vulnerability rather than overt neurodegeneration. These changes may sensitize the affected neurons to further dysfunction and death when challenged by additional pathological insults. In line with this hypothesis, our previous work showed that induction of tau pathology in the same mouse model leads to pronounced hippocampal and cortical atrophy (Palomares et al., 2025), whereas amyloid pathology has a comparatively milder and more regionally restricted impact (Suelves et al., 2023).

Our findings further demonstrate that mitochondrial impairment precedes energy failure during telomere-driven brain senescence. Although G3Terc^-/-^ mice exhibited reduced mitochondrial respiration as early as 5 months of age, a decline in energy charge was only observed at 9 months. These observations suggest that compensatory mechanisms in the adult mouse brain may initially preserve cellular energy homeostasis in the brain, masking the consequences of impaired OXPHOS during early senescence. Although OXPHOS is the principal source of ATP in neurons, ATP levels can be transiently maintained through metabolic adaptations, such as enhanced glycolysis and increased utilization of alternative energy substrates, including astrocyte-derived lactate (Area-Gomez, Guardia-Laguarta, Schon, & Przedborski, 2019). The progressive loss of these compensatory mechanisms with age may eventually result in bioenergetic failure and increased neuronal vulnerability. Future investigations should define these compensatory mechanisms and establish how they influence the progression of brain senescence.

Previous studies have reported that peripheral tissues from Terc knockout mice (e.g., liver, heart) display a significant decrease in mtDNA content, likely due to impaired mitochondrial biogenesis (Sahin et al., 2011). Surprisingly, our data revealed that brain tissue from these mice does not present changes in mtDNA content (indicative of total mitochondrial mass) or in overall ETC complex integrity, while still displaying altered OXPHOS activity. Differences compared with previously published data may reflect tissue-specific metabolic responses to telomere dysfunction. In contrast, G3Terc^-/-^ primary neurons derived from postnatal mouse pups and cultured for two weeks in vitro display a significant increase in mitochondrial volume. A similar increase in vitro has previously been reported in hepatocytes with telomere dysfunction induced by TRF2 deletion (Sullivan et al., 2023). This mitochondrial increase may represent an adaptive response aimed at compensating mitochondrial dysfunction to maintain ATP production and cellular energy homeostasis. This aligns with our observation that respiratory capacity is not impaired in these neurons. Mitochondrial functional defects may only become apparent in G3Terc^-/-^ neurons after extended time in vitro or following exposure to additional pathological stressors. Alternatively, the accumulation of mitochondria could also reflect a defective mitochondrial clearance through mitophagy, since alterations in autophagy have previously been reported by our group in G3Terc^-/-^ primary neurons (Suelves et al., 2023).

In conclusion, our study revealed mitochondrial dysfunction as one of the key pathways downregulated in a mouse model of telomere-induced senescence and suggests that this impairment contributes to the selective vulnerability of specific neuronal populations. Together, our findings provide mechanistic insight into pathological brain aging and identify cellular pathways that may represent attractive targets for interventions aimed at preserving brain function with age.

## 4 Materials and Methods

### 4.1 Animals

Terc knockout mice (Blasco et al., 1997) (Strain #004132, The Jackson Laboratory), carrying a germline deletion for the telomerase RNA subunit Terc, were bred. All mice were maintained on a C57BL/6 genetic background. Terc^-/-^ mice were intercrossed to obtain different generations (G) of mice, up to G3. Genotyping was performed by polymerase chain reaction (PCR) using DNA extracted from ear tissue. Animals were housed on a 12 h light/12 h dark cycle in standard animal care facilities with access to food and water ad libitum. Age-matched male and female mice were used for all analyses.

### 4.2 Primary neuronal cultures

Primary cultures of neurons were prepared from postnatal day 0 (P0) mouse pups as previously described (Suelves et al., 2023). Briefly, brains were collected and placed in ice-cold HBSS supplemented with 0.2% glucose. After removal of the meninges, cortical and hippocampal tissues were dissected and incubated at 37°C for 3 minutes (min) in HBSS/0.2% glucose containing 10 mg/mL trypsin and 1 mg/mL deoxyribonuclease I (DNase I) (Worthington Biochemical). Tissues were then mechanically dissociated in Neurobasal™ medium supplemented with 0.5 mg/mL DNase I by gentle trituration using a glass pipette. Dissociation was repeated with a flame-narrowed glass pipette and samples were allowed to sediment for 5 min. Supernatants containing isolated neurons were carefully placed on top of fetal bovine serum (FBS) and centrifuged at 1000× g for 10 min. Pellets were resuspended in Neurobasal™ medium supplemented with 1 mM L-glutamine (ThermoFisher Scientific), 2% B-27™ Supplement (ThermoFisher Scientific) and 0.1% penicillin-streptomycin (ThermoFisher Scientific). Cultures were produced individually from each pup. Cells were plated in 24-,12-, 6-well or 96-well plates pre-coated with 10 μg/mL of poly-L-lysine (Sigma-Aldrich) and maintained at 37°C in a humidified atmosphere containing 5% CO2. Neurons were collected at DIV14 or DIV21 and processed for subsequent analyses.

To evaluate activity-dependent transcription, neurons were treated at DIV14 with either vehicle (DMSO diluted 1:2000 in culture medium) or Bicuculline (50 μM, from a stock at 100 mM diluted in DMSO) for 2 h before collection for protein analysis.

### 4.3 Primary astrocyte cultures

Primary astrocyte cultures were obtained from mouse pups aged 2 days as previously described (Contino et al., 2020). Briefly, cortices were isolated on ice-cold HBSS and dissociated by sequentially using a glass pipette and a flame-narrowed glass pipette. Samples were centrifuged 1000×g for 5 min. Pellets were resuspended in HBSS and centrifuged at 1,700 × g for 20 min on a 30% Percoll gradient. Astrocytes were collected from the interphase, washed in HBSS and centrifuged for 5 min at 1,500 × g. Pellets were resuspended and plated in DMEM-glutaMAX medium (Thermo Fisher Scientific) supplemented with 10% FBS (Biowest), 1% penicillin-streptomycin (ThermoFisher Scientific), 50 µg/mL L-proline (ThermoFisher Scientific), and 2.5 µg/mL amphotericin B (ThermoFisher Scientific). Cells were left to proliferate in flasks for 15 days at 37°C and 5% CO_2_, and media were changed every 4–5 days. After 15 days, astrocytes were plated and further cultured in DMEM-glutaMAX with 10% FBS. Two days later, differentiation was induced by reducing the concentration of FBS to 3% for 7 days prior to cell collection.

### 4.4 Telomere length analysis by quantitative PCR

Brain tissue was digested with proteinase K (100 μg/mL) in lysis buffer (100 mM NaCl, 100 mM Tris-HCl [pH 8.0], 1 mM EDTA [pH 8.0] and 1% SDS) at 55 °C overnight. DNA was purified using the phenol-chloroform extraction method. To measure the length of mouse telomeres, a quantitative PCR method was used as previously described (Suelves et al., 2023). Briefly, two separate quantitative PCR reactions were performed, one with telomere-specific primers and the other using primers for the single-copy gene *36B4* (Sigma-Aldrich, Table S4). Relative telomere length (expressed as T/S ratio) was calculated by comparing telomere amplification (T) with that of the single-copy gene (S) using the 2^−ΔΔCT^ method. Results were then normalized (fold change) to the control condition.

### 4.5 Western blot

Mouse brain tissues were homogenized by sonication in ice-cold lysis buffer containing 20 mM Tris base (pH 8.0), 150 mM NaCl, 1% NP-40, 10% glycerol and supplemented with protease and phosphatase inhibitor cocktails (Roche). Samples were centrifuged at 16,000 x g for 20 min at 4°C, and the resulting supernatants were collected and stored at -80°C. Isolated mitochondrial fractions were processed using the same procedure.

For the analysis of cellular lysates, cells were collected in ice-cold lysis buffer containing 50 mM Tris base (pH 7.5), 150 mM NaCl, 2 mM EDTA, 1% NP-40 and supplemented with protease and phosphatase inhibitor cocktails (Roche). Samples were sonicated and centrifuged at 10,000 g for 10 min at 4°C, and the supernatants collected and kept at -80°C.

Total protein concentrations were determined using the Pierce™ BCA Protein assay (Invitrogen, Thermo Fisher Scientific). Protein extracts (10-30 µg) were mixed with NuPAGE™ LDS Sample Buffer (Invitrogen, Thermo Fisher Scientific) and 50 mM dithiothreitol (DTT), and heated for 10 min at 70°C. Samples were resolved by SDS-PAGE electrophoresis on precast NuPAGE™ 4-12% Bis-Tris gels (Thermo Fisher Scientific) and MES-SDS running buffer (Thermo Fisher Scientific). SeeBlue™ Plus2 pre-stained (Thermo Fisher Scientific) was used as standard. Samples were then transferred for 2 h at 30 V with NuPAGE™ transfer buffer (Thermo Fisher Scientific) onto 0.45 μm nitrocellulose membranes (Thermo Fisher Scientific). After incubation for 30 min in blocking buffer containing 5% non-fat powdered milk in PBS/0.1% Tween®20 (PBS-T), membranes were blotted overnight at 4°C with primary antibodies (Table S5) diluted in PBS-T. Horseradish-peroxidase (HRP)-conjugated species-specific secondary antibodies (Sigma-Aldrich) were used to bind primary antibodies prior to ECL detection. ImageJ software (National Institutes of Health) was used to quantify protein band intensity, normalized to the protein band intensity of Actin, Gapdh, or TOM20 (for mitochondrial fractions).

### 4.6 Immunofluorescence in cultured cells

Primary neurons were seeded in 24-well plates at a density of 200,000 cells per well. At 21DIV, cells were carefully washed with PBS and fixed with PBS/4% paraformaldehyde for 10 min. Cells were rinsed three times with PBS and then permeabilized with a solution of PBS/0.3% Triton for 30 min and non-specific sites were blocked with PBS/0.3% Triton/5% FBS/3% NGS for 30 min. Primary antibodies (Table S5) diluted in the blocking solution were incubated overnight at 4°C. After three washes with PBS, cells were incubated with DAPI (Sigma-Aldrich, dilution 1:10,000) and appropriate Alexa Fluor-conjugated secondary antibodies diluted in the blocking solution. Finally, samples were consecutively washed in PBS and water, and mounted with Mowiol.

Images were acquired with an EVOS™ FL Auto fluorescence microscope. Blind manual quantification of γH2AX foci was performed with ImageJ software. REST signal mean density was analyzed only in areas with MAP2-positive signal with ImageJ. As negative controls, samples were processed as described in the absence of primary antibody and no signal was detected.

### 4.7 Electrochemiluminescence immunoassay (ECLIA) for SASP quantification

SASP components (IL-1β, IL-6, CXCL1 and TNF-α) were analyzed in conditioned medium from DIV14 primary mouse neurons using the Meso Scale Discovery V-PLEX Custom Mouse Biomarkers Proinflammatory Panel1 assay kit (K152A0H-1, MSD) according to the manufacturer’s instructions. Culture media samples were concentrated 10-fold by lyophilization and analyzed without further dilution. IL-1β levels did not reach the minimum detection threshold and were therefore excluded from further analyses. Cytokine levels were normalized to the total protein content of the corresponding cell lysates, as determined using the Pierce™ BCA Protein Assay Kit (Invitrogen, Thermo Fisher Scientific).

### 4.8 RNA-Sequencing

#### Sample preparation

Hippocampal tissues were collected from 5-month-old WT and G3Terc^-/-^ mice (n = 4 animals per genotype). Tissues were homogenized in TriPure™ Isolation Reagent (Roche) and total RNA was extracted according to the manufacturer’s protocol. RNA was further purified using the ReliaPrep RNA Tissue Miniprep System (Promega). Concentration and purity were assessed spectrophotometrically (BioSpec-nano; Shimadzu Biotech) and RNA integrity was evaluated using the Agilent RNA 6000 Nano Kit on an Agilent 2100 Bioanalyzer (Agilent Technologies); only samples with RNA Integrity Number (RIN) ≥ 8.0 were included. Samples (200 ng/µl) were sent to Macrogen Europe B.V. for library preparation (TruSeq Stranded mRNA) and paired-end sequencing (NovaSeq 6000 platform, 150-bp paired-end reads; ∼60 million reads per sample). As the library preparation method is based on poly(A) selection, and Terc is a non-polyadenylated non-coding RNA, Terc transcript levels could not be assessed by RNA-seq. The Terc knockout genotype was confirmed independently by PCR using DNA extracted from ear tissue.

#### Bioinformatic analysis

Raw FASTQ files were quality-checked with FastQC (v0.11.8) and adapter sequences and low-quality bases were trimmed using Trimmomatic (v0.39) (Bolger, Lohse, & Usadel, 2014). Trimmed reads were aligned to the mouse reference genome (GRCm38/mm10) using HISAT2 (v2.2.1) (Kim, Langmead, & Salzberg, 2015). Gene-level read counts were quantified from the aligned BAM files using featureCounts (Subread v2.0.3) (Liao, Smyth, & Shi, 2014) with the Ensembl annotation file Mus_musculus.GRCm38.94.gtf. Only uniquely mapping reads were retained. Differential gene expression analysis was performed with DESeq2 (v1.50.2) (Love, Huber, & Anders, 2014). To evaluate the potential influence of sex on gene expression, an initial model including sex as a covariate was tested. As the results were highly comparable to those obtained with a model based solely on genotype, all subsequent analyses were conducted using the simpler design formula ∼ genotype. p-values were adjusted for multiple comparisons using the Benjamini-Hochberg procedure. Genes with an adjusted p-value (padj) < 0.05 were considered as differentially expressed.

### 4.9 Quantitative RT-PCR

Total RNA was extracted using TriPure™ Isolation reagent (Roche) according to the manufacturer’s protocol. RNA was resuspended in DEPC-treated water and concentration and purity were assessed spectrophotometrically (BioSpec-nano; Shimadzu Biotech). Reverse transcription (RT) was carried out with the iScript cDNA synthesis kit (Bio-Rad Laboratories) using 1 μg of total RNA in a total volume reaction of 20 μL. To obtain a negative control, the iScript reverse transcriptase was omitted in the cDNA synthesis step. Quantitative PCR was performed using the appropriate primers (Sigma-Aldrich, Table S4) and the GoTaq® qPCR Master Mix (Promega), following manufacturer’s instructions. Relative quantification was calculated by the 2^−ΔΔCT^ method using *Gapdh and Tbp* as housekeeping genes. Results were then normalized (fold change) to the control condition.

### 4.10 Mass Spectrometry

#### Protein Sample Preparation

Sample preparation and analysis were performed in the Mass Spectrometry Facility from the University of Namur. Hippocampal tissues were collected from 5-month-old WT and G3Terc^-/-^ mice (n = 8 animals per genotype). Tissue samples were homogenized in 200 µL urea lysis buffer (8 M urea, 2% SDS, 1 mM EDTA, 50 mM Tris-HCl, pH 8.0) supplemented with protease and phosphatase inhibitors. Homogenization employed zirconium beads and a FastPrep-24 (MP Biomedicals) device (3× 20 s cycles at 6.5 m/s with 2-min cooling intervals at 4°C). Samples were sonicated using a Bioruptor (Diagenode)(5 × 30 s bursts, 60 s intervals on ice), then centrifuged at 20,000 x g for 5 min at 4°C. The clarified supernatant was collected, and protein concentration determined by Pierce 660 nm assay (ThermoFisher). Protein aliquots were stored at -80°C prior to proteomic analysis.

#### Protein Digestion

Samples were treated using Filter-aided sample preparation (FASP) using the following protocol. To first wash the filters, 100 µl of 1% formic acid was added to each Microcon 30 filter unit (Millipore), followed by centrifugation at 20,000 x g for 15 min. For each sample, 40 µg of protein was brought to a final volume of 150 µL using 8 M urea buffer (urea 8 M in buffer Tris 0.1 M at pH 8,5) and was loaded into a column and centrifuged at 20,000 x g for 15 min. The filtrate was discarded and the columns were washed three times by adding 200 µL of urea buffer followed by a centrifugation at 20,000 x g for 15 min. For the reduction step, 100 µL of 8 mM dithiothreitol (DTT) was added to the columns and mixed for 1 min at 400 rpm with a thermomixer before an incubation of 15 min at 24°C. Samples were then centrifuged at 20,000 x g for 15 min, the filtrate was discarded and the filter was rinsed by adding 100 µL of urea buffer, followed by another centrifugation at 20,000 x g for 15 min. An alkylation step was performed by adding 100 µL of 50 mM iodoacetamide (IAA), in urea buffer) to each column, followed by mixing at 400 rpm for 1 min in the dark. The samples were then incubated for 20 min in the dark and centrifuged at 20,000 x g for 10 min. To remove the excess of IAA, 100 µL of urea buffer was added and the samples were centrifugated at 20,000 x g for 15 min. To quench residual IAA, 100 µL of DTT was placed on the column, mixed for 1 min at 400 rpm and incubated for 15 min at 24 °C before centrifugation at 20,000 x g for 10 min. To remove the excess of DTT, 100 µL of urea buffer was added to the column and centrifuged at 20,000 x g for 15 min. The filtrate was discarded, and the column was washed three times by adding 100 µL of sodium bicarbonate buffer 50 mM (ABC) in ultrapure water, followed by a centrifugation at 20,000 x g for 10 min. The last 100 µL of buffer was left at the bottom of the filter unit to avoid any evaporation in the column. The protein digestion process was performed by adding 80 µL of mass spectrometry-grade trypsin (1/50 in ABC buffer) to the column, followed by mixing at 400 rpm for 1 min before an incubation overnight at 24°C in a water-saturated environment. The Microcon columns were placed into protein LoBind tubes (Eppendorf) of 1.5 mL and centrifuged at 20,000 x g for 10 min. 40 µL of ABC buffer was placed on the column before centrifugation at 20,000 x g for 10 min. 10% Trifluoroacetic acid (TFA) in ultrapure water was added to the content of the LoBind tube to obtain a final concentration of 0.2 % TFA. The samples were dried in a SpeedVac to a final volume of 20 µL and transferred into injection vials.

#### LC-Mass spectrometry

The digest was analyzed using nano-LC-ESI-MS/MS tims TOF Pro (Bruker, Billerica, MA, USA) coupled with an UHPLC nanoElute (Bruker). The different samples were analysed with a gradient of 90 min. Peptides were separated by nanoUHPLC (nanoElute, Bruker) on a 75 μm ID, 25 cm C18 column with integrated CaptiveSpray insert (Aurora, ionopticks, Melbourne) at a flow rate of 200 nl/min, at 50°C. LC mobile phases A was water with 0.1% formic acid (v/v) and B, ACN with formic acid 0.1% (v/v). Samples were loaded directly on the analytical column at a constant pressure of 800 bar. The digest (1 µl) was injected, and the organic content of the mobile phase was increased linearly from 2% B to 15 % in 55 min, from 15 % B to 24% in 27 min, from 24% to 35% B in 8 min. Data acquisition on the tims TOF Pro was performed using Hystar 6.1 and timsControl 2.0. tims TOF Pro data were acquired using 160 ms TIMS accumulation time, mobility (1/K0) range from 0.75 to 1.42 Vs/cm². Mass-spectrometric analysis was carried out using the parallel accumulation serial fragmentation (PASEF) (Meier et al., 2018) acquisition method, with one MS spectrum followed by six PASEF MS/MS spectra per total cycle of 1.16 s.

#### Mass spectrometry data processing

LC/MS/MS raw data were processed with the MaxQuant software (version 2.4.9.0, Max Planck Institute of Biochemistry, Munich, Germany) using the Andromeda module with default search settings. Spectra were searched against a Mus musculus protein database including isoforms and common contaminants (UniProt Proteome ID: UP000000589, 63,642 sequences). The Trypsin/P enzyme parameter was selected with one possible missed cleavage. Carbamidomethylation of cysteines was set as a fixed modification, while methionine oxidation and acetylation (protein N-term) were selected as variable modifications. For protein validation, a maximum false discovery rate of 1% at peptide and protein levels was used based on a decoy search. The mass tolerance was set to 20 ppm for peptide masses during the first search, and 10 ppm during the main search. A minimum of one unique peptide was required for identification. Matching between runs was performed using a 0.8 min match time window, a 0.05 1/K_0_ ion mobility window and a 20 min alignment time window.

#### Bioinformatic analysis

Differential protein abundance analysis was carried out MaxQuant-derived LFQ intensities using the QFeatures (v1.20.0) (Gatto & Vanderaa, 2024) and msqrob2 (v1.18.0) (Sticker, Goeminne, Martens, & Clement, 2020) packages. Statistical modeling was performed using a linear model with genotype as the main factor of interest and batch included as a covariate to correct for technical variation. Gene Set Enrichment Analysis (GSEA) was performed using clusterProfiler v4.20.0 (Yu, Wang, Han, & He, 2012) on genesets from the molecular Signature Database (MSigDB) downloaded using msigdbr package v26.1.0.

To assess the concordance between transcriptomic and proteomic differential analyses, genes and proteins quantified in both datasets were matched based on their gene symbols. The moderated *t*-statistics obtained from the DESeq2 (RNA-seq) and msqrob2 (proteomics) differential analyses were then compared by calculating the Pearson correlation coefficient across all matched features.

### 4.11 Mitochondrial isolation and Electron flow assay

Hippocampal and cortical tissues were freshly collected from adult mice sacrificed by cervical dislocation. Tissues were rapidly weighed and homogenized with a Dounce glass homogenizer with 50 μL/mg tissue of BSA-free mitochondrial assay solution (MAS) containing 2 mM HEPES, 70 mM sucrose, 220 mM mannitol, 10 mM KH₂PO₄, 5 mM MgCl₂, and 1 mM EGTA (pH 7.2). The initial homogenate (TH1) was centrifuged at 800 x g for 10 min at 4°C and the resulting supernatants (TH2) were collected and centrifuged once again at 8000 x g for 15 min at 4°C. The resulting supernatants (CF) contained the cytosolic fraction, while the pellets, containing the crude mitochondrial fraction, were washed once with BSA-free MAS buffer and resuspended in 3 μL of BSA-free MAS buffer per mg of tissue (MF). Total protein concentration was determined using a standard BCA assay. Western blot analyses were performed on the different fractions (TH1, TH2, CF, and MF) to confirm the enrichment of intact mitochondria in the final mitochondrial fraction (Figure S8). Of note, as this preparation consisted of a crude mitochondrial isolation, some cytosolic and nuclear markers were present in the mitochondrial fraction.

To measure the activity of the different complexes of the electron transport chain, the Agilent Seahorse XF Electron Flow Assay was performed on isolated mitochondria from adult mice, following Agilent’s recommendations and as previously described (Corbet et al., 2016; de Mey et al., 2018). Freshly isolated mitochondria were diluted in MAS supplemented with 0.2% (w/v) fatty acid-free BSA, and seeded (4 μg in 25 μL per well) in XFe96 Seahorse plates pre-coated with 10 μg/mL poly-L-lysine (Sigma-Aldrich), followed by centrifugation at 2000 × g for 20 min at 4°C. The mitochondria were viewed briefly under a microscope to ensure consistent adherence to the well. 150 µL of pre-warmed MAS supplemented with fatty-acid-free BSA 0.2% and with the mitochondrial substrates pyruvate (10 mM), malate (2.5 mM), and ADP (2 mM), were added to each well containing the isolated mitochondria. Determination of complexes I-IV-dependent respiration was accomplished by the sequential injection of rotenone (1 μM; complex I inhibitor), succinate (10 mM; complex II substrate), antimycin A (4 μM; complex III inhibitor) and ascorbate/TMPD (N,N,N′,N′-tetramethyl p-phenylenediamine) (10 mM/100 μM; electron donors to complex IV). Following completion of the electron flow assay, total protein concentration was determined using a Bradford assay for normalization purposes.

The individual activities of ETC Complex I-IV can be deduced by way of subtraction of the OCR values as previously described (de Mey et al., 2018). Briefly, the activity of complex I was measured by subtracting the OCR after the injection of rotenone from the OCR at baseline ETC activity (measured when mitochondria are incubated with pyruvate and malate). Complex II activity was calculated by the subtraction of the OCR after the injection of rotenone from the OCR after the injection of succinate. Complex III activity was obtained by subtracting the OCR after antimycin A injection from the OCR after the injection of succinate. Finally, the activity of complex IV was measured by subtraction of the OCR after antimycin A injection from the OCR values after ascorbate and TMPD injection.

### 4.12 Long-extension PCR for mtDNA damage evaluation

Mitochondrial DNA damage was assessed by long-range PCR amplification of DNA extracts. Briefly, mouse brain tissue was digested with proteinase K (100 μg/mL) in lysis buffer (100 mM NaCl, 100 mM Tris–HCl [pH 8.0], 1 mM EDTA [pH 8.0] and 1% SDS) at 55°C overnight. DNA was purified using the phenol-chloroform extraction method. Mitochondrial DNA was amplified from 2 ng of total DNA with the appropriate forward and reverse primers (Eurogentec, Table S4) targeting a 3.5 kb fragment within the mitochondrial minor arc, using the KAPA2 Robust Hotstart PCR kit (Roche) and following previously described PCR conditions (Matic et al., 2018). The integrity of the resulting mitochondrial amplicons was assessed by electrophoresis in 1% agarose gels and visualized by Midori Green Advance (Nippon Genetics) staining.

### 4.13 Quantification of total mitochondrial DNA content

Relative mitochondrial DNA (mtDNA) content was quantified by qPCR from isolated total DNA as previously described (Contino et al., 2020). Briefly, brain tissue or primary neurons were digested with proteinase K (100 μg/ml) in lysis buffer [100 mM NaCl, 100 mM Tris–HCl (pH 8.0), 1 mM EDTA, (pH 8.0) and 1% (w/v) SDS] at 55 °C o/n and genomic DNA was purified with phenol/chloroform method. Quantitative PCR was performed using different primer pairs to detect the mitochondrial genes *Cyt-b* and *Nd1*, and the nuclear gene *H19* (Table S4). The relative mitochondrial DNA content was calculated by the 2^−ΔΔCT^ method. Results were then normalized (fold change) to the control condition.

### 4.14 HPLC nucleotide measurement

Mice were sacrificed by cervical dislocation followed by decapitation and rapidly dissected in an ice-cold extraction solution containing 0.1 M HClO₄ and 40% methanol to inhibit enzymatic activity and preserve ATP levels. Hippocampal and cortical brain tissues were collected as fast as possible and snap-frozen in liquid nitrogen before being placed in dry ice. Once samples were weighed, they were homogenized in cold extraction solution using an Ultra-Turrax apparatus with a medium-sized head. After lysis, the samples were centrifuged at 15,000 x g for 5 min at 4°C. The resulting supernatants were collected and neutralized with 1.1M ammonium phosphate before being dried using a SpeedVac. Dried samples were resuspended in HPLC-grade water and vortexed for 5 min, sonicated for 2 min, and vortexed once again for 2 min to ensure proper resuspension of the pellets. Finally, the samples were centrifuged at 3,000 x g for 2 min at room temperature, and the resulting supernatants were transferred to HPLC vials for analysis.

An AtlantisTM T3 (3 μm, 3x100 mm) C18 column (Waters) specifically designed for the separation of purine nucleotides was chosen as our stationary phase. The mobile phase consisted of a 0.1M ammonium phosphate buffer (pH 6) with a gradient of methanol increasing to 50%. UV detection was employed at multiple wavelengths, with 262 nm selected as the optimal wavelength for observing the highest peaks of ATP, ADP, and AMP. Identification of individual peaks was achieved by combining retention times with UV absorbance profiles. Retention times were determined using external standards with known concentrations, while quantification was performed based on the area under the UV peaks. These areas were converted to absolute quantities using the standard curves. Data analysis, including calculation of the area under the curve, was carried out using OpenLab CDS software (Agilent). Final nucleotide measurements were normalized to the tissue weight. The energy charge (EC) was calculated using the following equation:

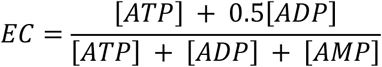

### 4.15 Evaluation of ROS levels in primary neurons

Intracellular ROS levels were measured using CM-H_2_DCFDA (Invitrogen) as a fluorescent probe. Primary neurons were seeded in 96-well plates at a density of 50,000 cells per well. At 14 DIV, cells were incubated for 30 min at 37°C and 5% CO₂ in HBSS supplemented with 1 μM CM-H_2_DCFDA (Invitrogen, C6827) and 2 μM Hoechst stain (Thermo Fisher Scientific). After incubation, neurons were washed with prewarmed HBSS and fluorescence images were acquired at 10X magnification using an EVOS microscope. All images were acquired using identical acquisition settings. The mean fluorescence intensity of the thresholded CM- H2DCFDA signal, with the background subtracted, was quantified using ImageJ software.

Alternatively, fluorescence signals were quantified using a Victor X4 microplate reader (PerkinElmer) with appropriate excitation and emission settings for CM-H_2_DCFDA and Hoechst. To induce oxidative stress, parallel cultures were treated with 1 mM hydrogen peroxide (H₂O₂), a well-established ROS inducer, and used as a positive control.

### 4.16 Evaluation of ATP levels in primary neurons

Intracellular ATP levels were measured using the CellTiter-Glo® Luminescent Cell Viability Assay (Promega, Wisconsin, USA) according to the manufacturer’s instructions. Primary neurons were seeded in 96-well plates at a density of 50,000 cells per well. On the day of the experiment, neurons were incubated with 100 μL culture medium containing 2 μM of Hoechst dye for 30 min at room temperature. Hoechst fluorescence intensity was measured using a Victor X4 microplate reader (PerkinElmer) at 461 nm to determine total cell number. Subsequently, 100 μL of CellTiter-Glo reagent was added to each well, and plates were mixed for 2 min on an orbital shaker to induce cell lysis. Plates were then incubated for 10 min at room temperature to stabilize the luminescent signal before luminescence was measured using the Victor X4 microplate reader. ATP levels were normalized to total cell number, as determined by Hoechst fluorescence measured with the plate reader.

### 4.17 Evaluation of mitochondrial respiration in primary neurons

Mitochondrial respiration was measured in primary mouse neurons using the Agilent Seahorse XF Mito Stress Test assay. Cells were seeded in XFe96 Seahorse plates pre-coated with 10 μg/mL poly-L-lysine (Sigma-Aldrich), at a density of 60,000/well, and cultured in Neurobasal^TM^ medium enriched with 1 mM L-glutamine, 5 mM glucose, 2% B-27™ Supplement and 0.1% penicillin–streptomycin. At DIV14, the culture medium was replaced with pre-warmed Seahorse assay medium (DMEM base, Sigma-Aldrich) supplemented with 1.85 g/L NaCl, 3 mg/L phenol red, 10 mM D-glucose and 2 mM L-glutamine, adjusted to pH 7.35 ± 0.05. Cells were then incubated for 60 min at 37°C in a non-CO_2_ incubator prior to the assay.

The oxygen consumption rate (OCR; pmol O_2_/min) was measured under basal conditions (three measurement cycles consisting of 1 min 30 s mixing, 2 min waiting and 3 min reading) and following the sequential injection of mitochondrial inhibitors: oligomycin (2 μM; three cycles consisting of 1 min 20 s mixing, 1 min 20 s waiting and 4 min reading), FCCP (0.5 μM; three cycles consisting of 20 s mixing and 2 min reading), and rotenone/antimycin A (0.5 μM/0.5 μM; three cycles consisting of 1 min 20 s mixing, 1 min 20 s waiting and 2 min reading). The pH of all drug solutions was adjusted to 7.35 ± 0.05 prior to use.

OCR values were normalized to the total protein content of each well determined after completion of the assay by the Bradford assay kit (Bio-Rad Laboratories). Mitochondrial respiration parameters were calculated according to the manufacturer’s recommendations. Non-mitochondrial respiration was defined as the OCR measured after rotenone/antimycin A injection. Basal respiration was calculated as basal OCR minus non-mitochondrial respiration. Respiration linked to ATP production was calculated as the difference between basal OCR and OCR following oligomycin treatment. Maximal respiration was calculated as FCCP-stimulated OCR minus non-mitochondrial respiration. Spare respiratory capacity was calculated as maximal respiration minus basal respiration.

### 4.18 Cell viability assay

Cell viability in primary mouse neurons was assessed using the CellTiter 96® AQueous One Solution Cell Proliferation Assay (Promega), which is based on similar principles to the widely used MTS assay. Following manufacturer instructions, cells were seeded on 96-well plates at a density of 60,000/well. At DIV14, 20 μL of Celltiter solution were added per well. Absorbance at 490nm was read after 4 hours of incubation using the Victor X4 microplate reader. Levels were normalized to total cell number, as determined by Hoechst fluorescence measured with a Victor X4 microplate reader (PerkinElmer).

### 4.19 Immunofluorescence in brain sections

Mice were anesthetized with a combination of ketamine (100 mg/kg, Dechra) and medetomidine (1 mg/kg, Vetoquinol) diluted in sterile phosphate-buffered saline (PBS), followed by transcardial perfusion with ice-cold sterile PBS. The brains were removed and post-fixed by immersion in 4% paraformaldehyde in PBS for 24 h, cryoprotected in PBS containing 30% sucrose and 0.02% sodium azide, and subsequently frozen.

Coronal brain sections (30 μm thick) were obtained using a CryoStar™ NX50 Cryostat (Thermo Fisher Scientific) and stored at –20 °C in cryoprotectant solution (sodium phosphate buffer containing 30% ethylene glycol and 20% glycerol) until use. Free-floating sections were selected, rinsed three times in 1X PBS and blocked/permeabilized for 30 min at room temperature in blocking solution containing 5% BSA, 3% normal goat serum (Gibco) and 0.5% Triton X-100 in 1X PBS. Brain sections were then incubated overnight at 4 °C with primary antibodies (Table S5) diluted in blocking solution. After three washes with 1X PBS, sections were incubated in the dark for 1 h at room temperature with DAPI (1:10,000, Sigma-Aldrich) and the appropriate species-specific fluorophore-conjugated Alexa Fluor™ secondary antibodies diluted in blocking solution. Finally, sections were washed three times in 1X PBS and mounted on Superfrost™ glass slides, air dried and coverslipped using Mowiol mounting medium. Sections were stored at 4°C. Images were captured using a ZEISS Axioscan 7 Microscope Slide Scanner with a 20X objective lens. The density of NeuN-positive cells was quantified using the open-source software QuPath (Bankhead et al., 2017). Briefly, the “cell detection” function in the analysis menu was used to automatically count the number of cells within a pre-defined area. Detection thresholds were optimized and kept constant across all samples to ensure comparability.

### 4.20 Statistical analysis

Statistical analyses were carried out using the GraphPad Prism 10 software. Data normality was assessed using the Shapiro–Wilk test. Parametric tests were applied when data followed a normal distribution, whereas non-parametric tests were used otherwise. Comparisons between two groups were performed using either an unpaired Student’s t-test or a Mann– Whitney test, as appropriate. For comparisons involving more than two groups, a one-way ANOVA followed by the indicated post hoc test or a Kruskal-Wallis test was used. When assessing the effects of two independent variables (e.g., genotype and treatment), a two-way ANOVA followed by the indicated post hoc test was performed. Significance is indicated as: ns = non-significant, *P < 0.05, **P < 0.01, ***P < 0.001. The number of independent biological samples (i.e., animals) is indicated in figure legends with “n = ”.

## Author Contributions

NS and PKC conceived, designed and supervised the research project. NS, DP, JJ, SS, TI, AL, MD, SB, and MJ performed experiments and analyzed and interpreted data. MJ, CC, PR, LG, and PKC contributed to the methodology and helped revise the manuscript. NS and DP wrote the manuscript. All authors have read and approved the final version of the manuscript.

## Acknowledgements

We thank Caroline Bouzin and Aurélie Daumerie from the 2IP Imaging Platform (Institut de Recherche Expérimentale et Clinique – IREC, UCLouvain, Brussels, Belgium) for their assistance with microscopy imaging and analysis.

## Funding

DP and JJ were supported by a FRIA fellowship from the Belgian FRS-FNRS (Fonds de la recherche scientifique). NS was funded by a Chargé de Recherche postdoctoral fellowship from the FRS-FNRS. This work was supported by grants from the SAO-FRA Alzheimer Research Foundation (SAO-FRA 2018/0025), UCLouvain Action de Recherche Concertée (ARC21/26–114), Fondation Louvain, Queen Elisabeth Medical Foundation (FMRE AlzHex), F.R.S.-FNRS (FNRS J.0106.22) attributed to PKC and grants from the SAO-FRA Alzheimer Research Foundation (SAO-FRA 2020/0028, SAO-FRA 2022/0028) attributed to NS.

## Conflicts of Interest

The authors declare no conflicts of interest.

## Data Availability Statement

Most datasets generated and analyzed during this study are included in this published article and its supplementary material. Any additional information required to reanalyze the data reported in this paper is available from the corresponding author upon reasonable request.

RNA-sequencing data have been deposited to NCBI Gene Expression Omnibus (GEO) database and will be made publicly available upon acceptance. Proteomics data have been submitted to PRIDE and will be made publicly available upon acceptance. Accession numbers will be provided upon publication.

## Notes

### Competing Interest Statement

The authors have declared no competing interest.

