## Supplementary Figures for "Mitochondrial dysfunction as a hallmark of brain senescence in telomerase-deficient mice"

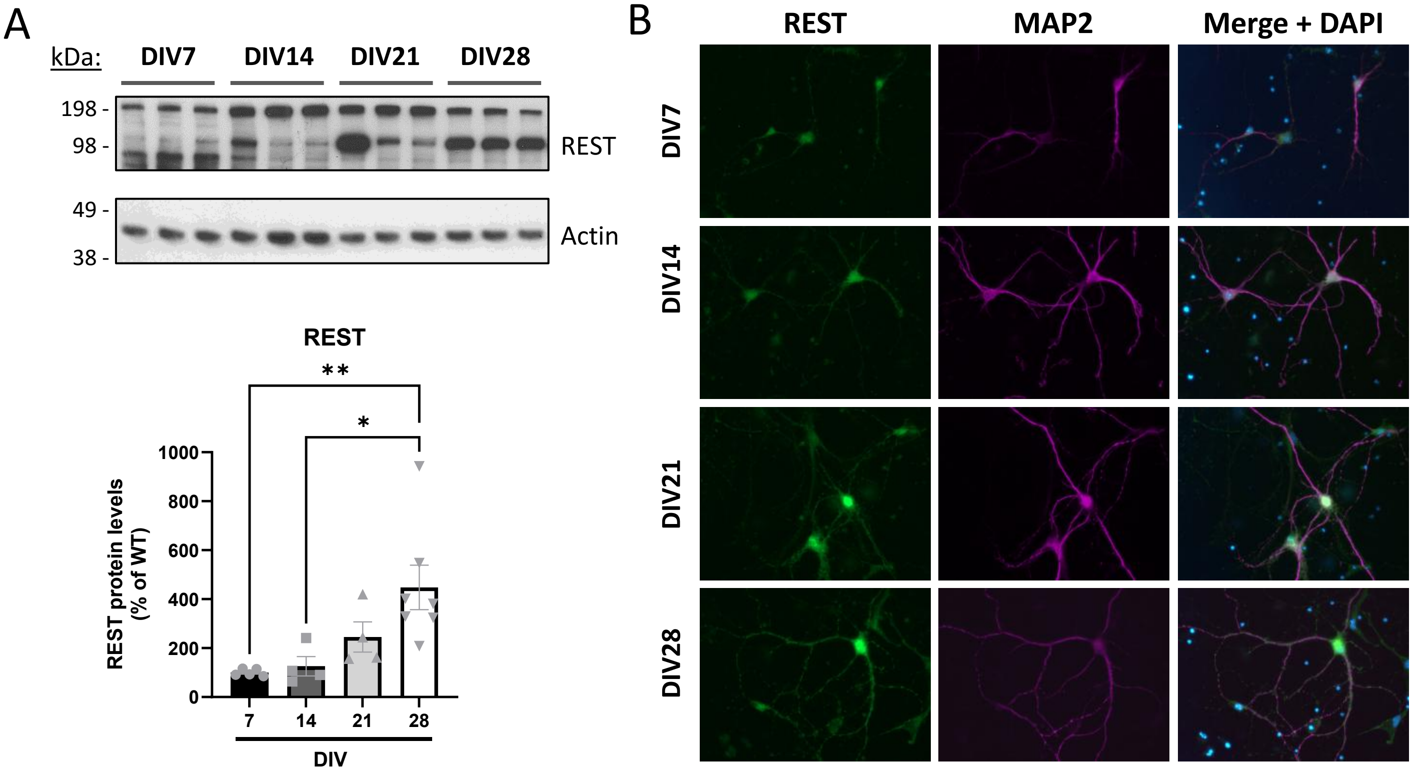


**Figure S1. REST expression increases over time in primary mouse neuronal cultures.** (A) Western blot analysis of REST protein levels in lysates from WT primary neuronal cultures collected at DIV7, DIV14, DIV21, and DIV28. REST protein levels were normalized to actin and expressed relative to DIV7, which was set to 100%. $^*P < 0.05$, $^{**}P < 0.01$ (One-way ANOVA with Tukey's post-hoc analysis, n = 4-7 cultures/time point). (B) Immunofluorescence analysis of REST (green) and MAP2 (magenta) in WT primary neuronal cultures collected at DIV7, DIV14, DIV21, and DIV28.

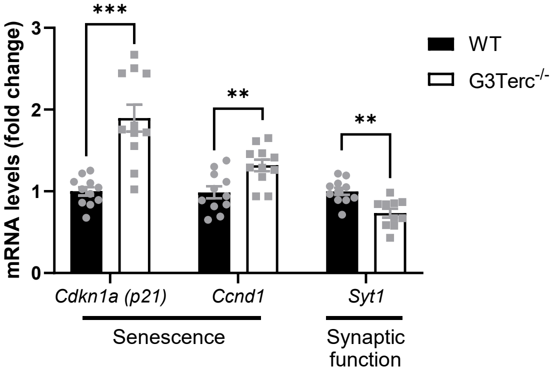


**Figure S2.** Validation of selected differentially expressed genes identified by transcriptomic analysis. mRNA levels of *Cdkn1a* (also known as *p21*), *Ccnd1*, and *Syt1* genes were measured by RT-qPCR in hippocampal extracts from 5-month-old WT and G3Terc$^{-/-}$ mice. *P < 0.05, **P < 0.01, ***P < 0.001 (two-tailed Student's t-test, n = 10 cultures/group).

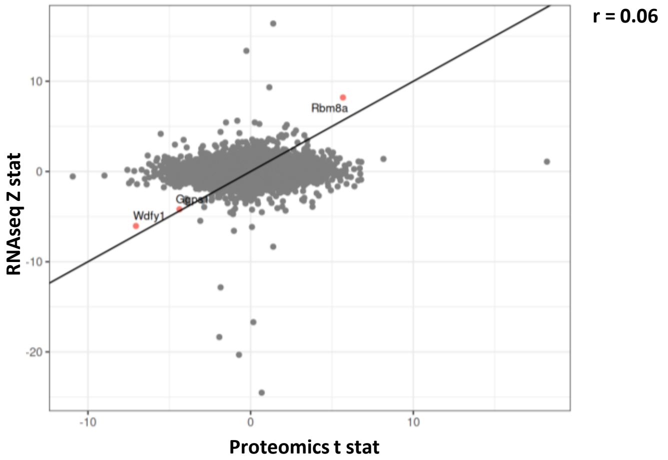


**Figure S3. Low correlation between the transcriptomic and proteomic data from our study.** Comparison of differential expression test statistics (proteomics t-statistic vs. RNA-seq z-statistic) from RNAseq and LC-MS/MS datasets obtained from hippocampi of 5-month-old WT and G3Terc$^{-/-}$ mice. Genes/proteins concordantly dysregulated at the mRNA and protein level are highlighted in red; all other datapoints are shown in grey. The Pearson's correlation coefficient (r) is 0.06.

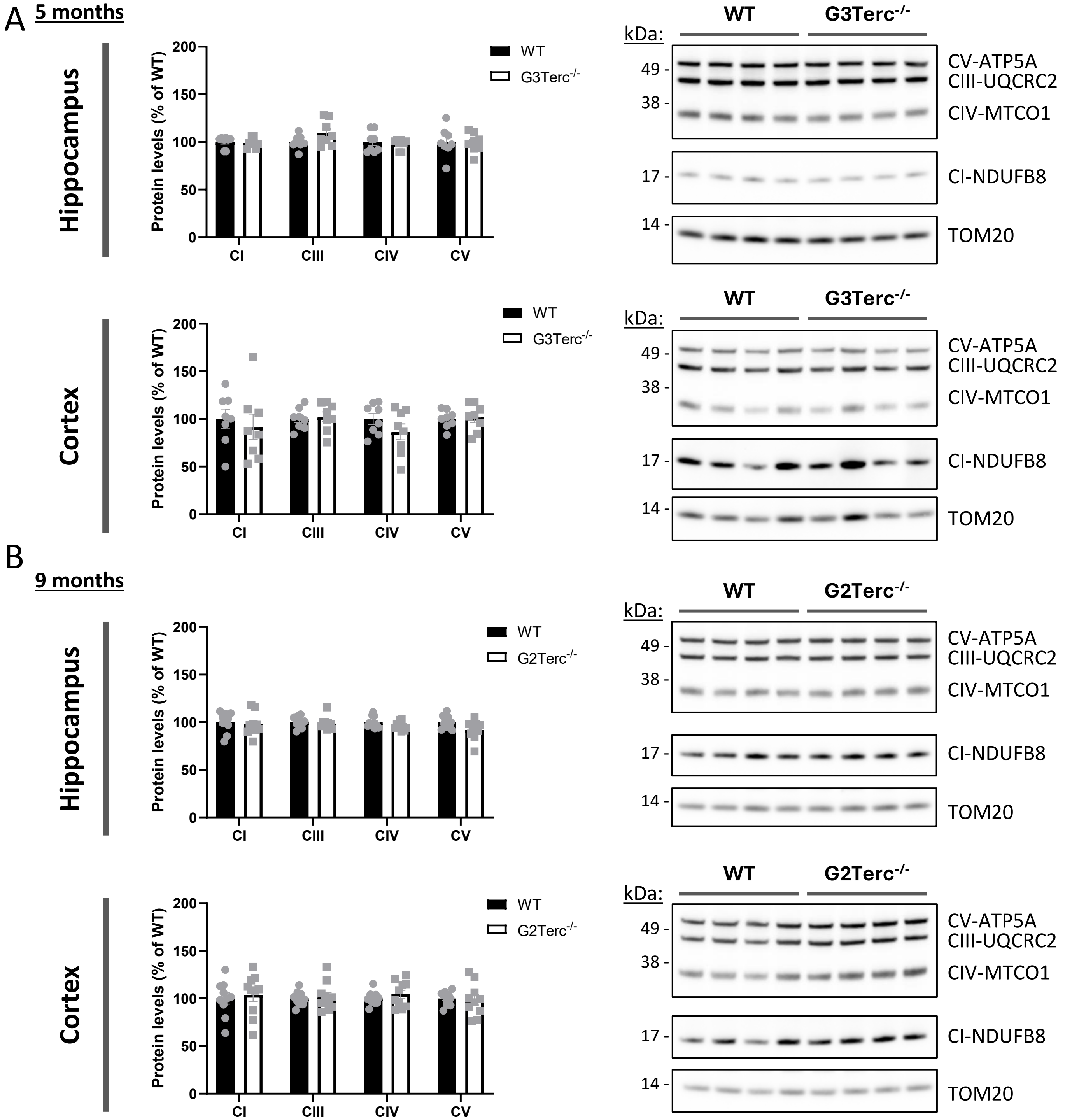


**Figure S4. Senescent brains show functional ETC impairments without gross alterations in ETC complex subunit levels.** (A, B). Western blot analysis of complex I, III, IV, and V main subunits in hippocampal and cortical mitochondrial extracts from 5-month-old WT and G3Terc$^{-/-}$ mice (A) or 9-month-old WT and G2Terc$^{-/-}$ mice (B). Complex II subunits were not included, as they could not be consistently detected with sufficient signal quality. TOM20 was used as a loading control, and levels in the WT group were normalized to 100%. Non-significant (two-tailed Student's t-test, n = 5 mice/group). All data are presented as mean ± SEM.

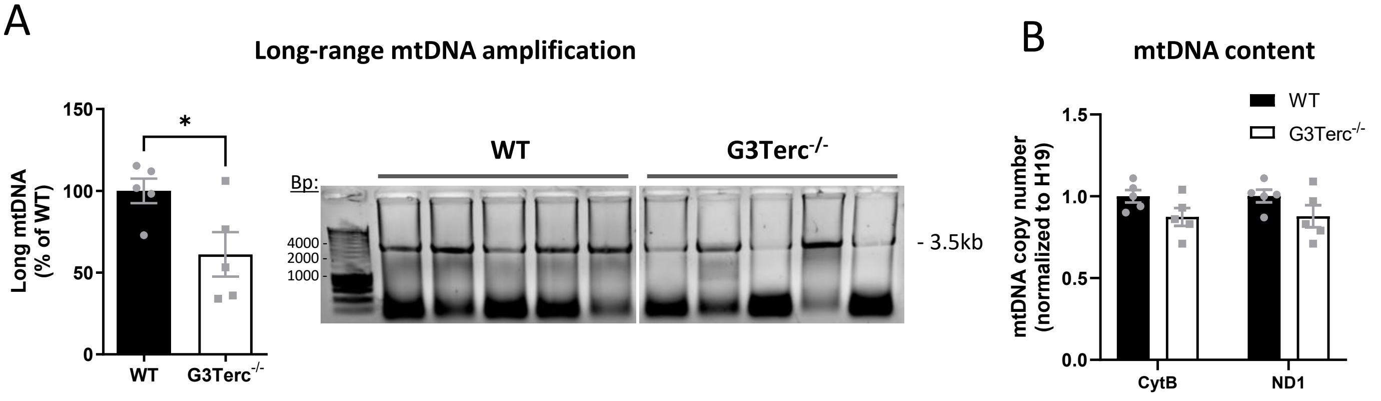


**Figure S5. Senescent brains display mtDNA damage with no changes in total mitochondrial DNA.** (A) PCR analysis of long mitochondrial DNA amplification in hippocampal extracts from WT and G3Terc$^{-/-}$ mice at 6 months of age. The relative amount of PCR product (3.5kb) was calculated and levels in the WT group were normalized to 100%. *P < 0.05 (two-tailed Student's t-test, n = 5 mice/group). (B) qPCR analysis of mitochondrial DNA content (*CytB* and *Nd1* genes) in hippocampal samples from WT and G3Terc$^{-/-}$ mice at 6 months of age. Non-significant (two-tailed Student's t-test, n = 5 mice/group). All data are presented as mean ± SEM.

A

| Gene set | Set size | ES | NES | p-value |
|---|---|---|---|---|
| Mitochondrial-encoded genes | 13 | -0.855183 | -2.231944 | 7.38395E-06 |

B

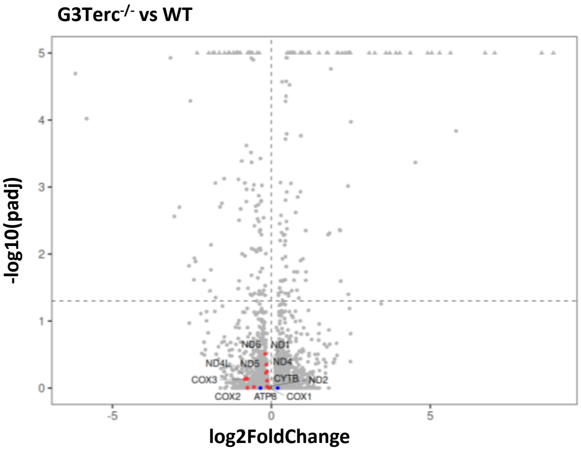


**Figure S6. Mitochondrially-encoded genes exhibit negative enrichment in senescent mouse brains.** (A) Targeted GSEA of Mitochondrial-encoded genes. (B) Volcano plot highlighting (in red) the Mitochondrial-encoded genes in the gene set evaluated. Data points for genes with p-adjusted values < $10^{-5}$ are shown as triangles at −log10(padj) = 5.

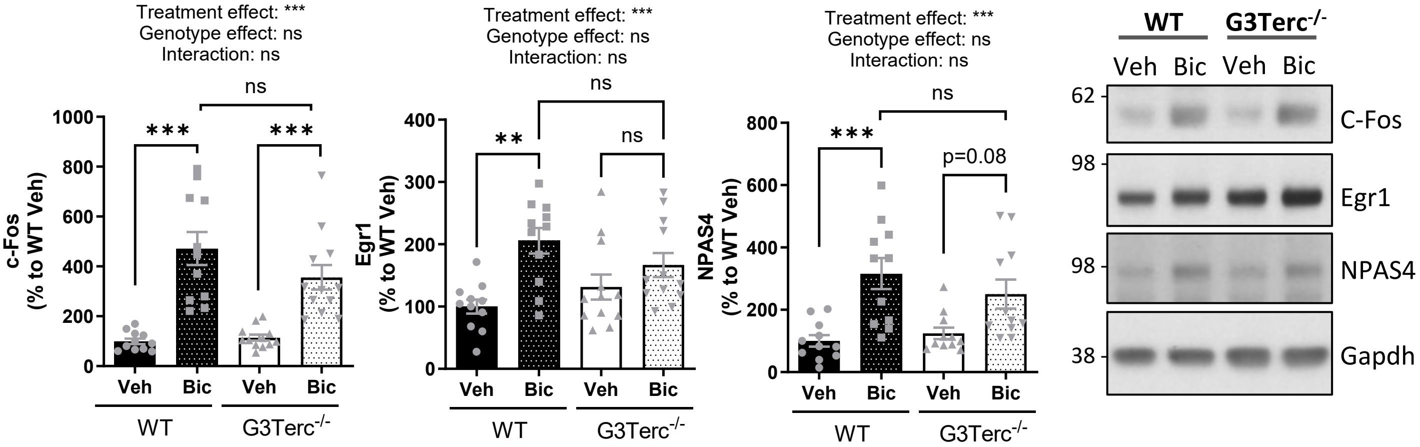


**Figure S7. Activity-dependent responses are not altered in G3Terc$^{-/-}$ primary neurons.** Western blot analysis of c-Fos, Egr1, and NPAS4 protein levels in DIV14 WT and G3Terc$^{-/-}$ primary neurons following stimulation with bicuculline (50 μM) or vehicle for 2 h. **P < 0.01, ***P < 0.001 (Two-way ANOVA with Tukey's post-hoc analysis, n = 11-12 cultures/group).

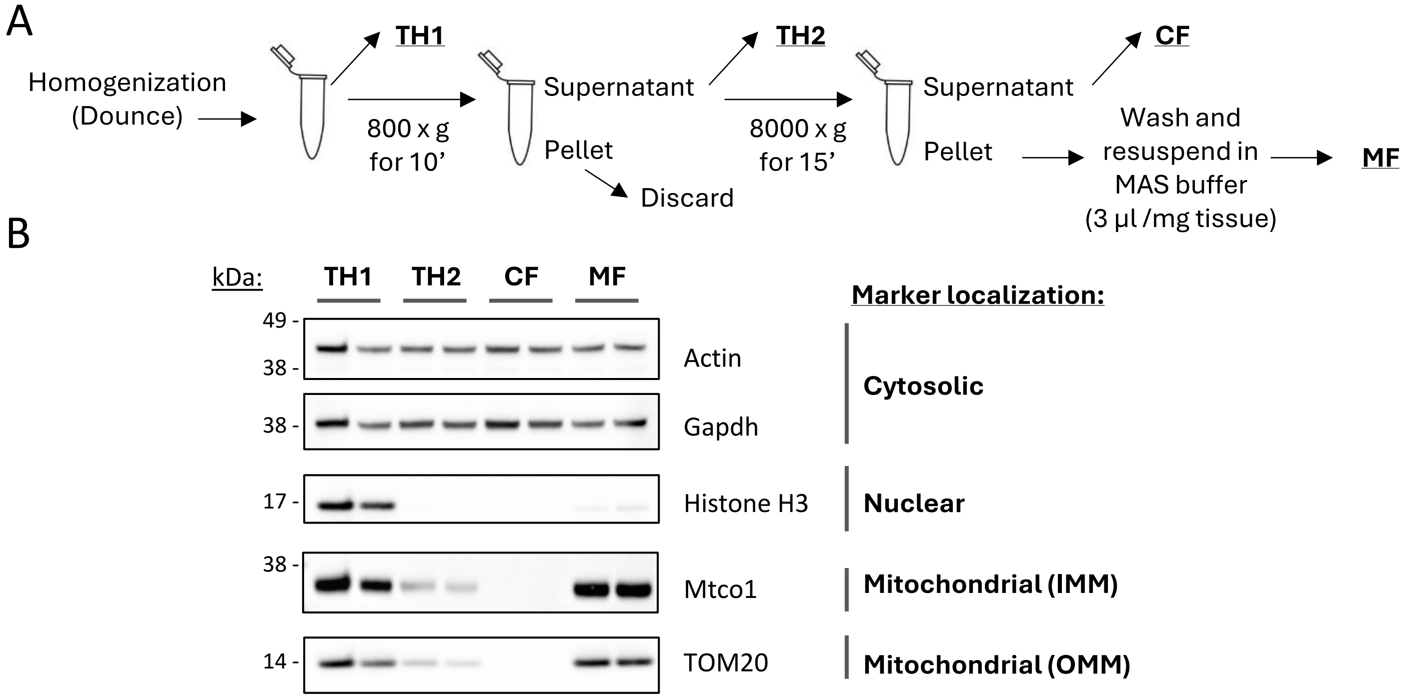


**Figure S8. Validation of the mitochondrial isolation protocol used for the Electron Flow assay.** (A) Schematic overview of the mitochondrial isolation procedure. Mouse brain mitochondria were isolated through sequential centrifugation steps, generating the following fractions: total homogenate 1 (TH1), obtained after the initial tissue homogenization prior to centrifugation; total homogenate 2 (TH2), corresponding to the supernatant collected after the first centrifugation step and largely depleted of nuclei; cytosolic fraction (CF); and mitochondrial fraction (MF). (B) Western blot analysis of the fractions collected during mitochondrial isolation to assess the purity and enrichment of the mitochondrial preparation. GAPDH and actin were used as cytosolic markers, Histone H3 as a nuclear marker, and TOM20 and MT-CO1 as markers of the outer mitochondrial membrane (OMM) and inner mitochondrial membrane (IMM), respectively.
